# A standardized framework resolves ambiguity in motor neuron loss across neurodegenerative diseases

**DOI:** 10.64898/2026.04.15.718647

**Authors:** Leonie Sowoidnich, Aaron L. Norman, Florian Gerstner, Josiane K. Siemund, Jannik M. Buettner, John G. Pagiazitis, Vanessa Dreilich, Konstantin Pilz, Dajun Tian, Charlotte J. Sumner, Angela Paradis, George Z. Mentis, Christian M. Simon

**Author notes:** Correspondence to: Christian M. Simon, Carl-Ludwig-Institute for Physiology, Leipzig University, Room E105, Liebigstraße 27, Leipzig, Saxony 04103, Germany. These authors contributed equally to this work (Leonie Sowoidnich, Aaron L. Norman). Employee of Biogen at the time of contributing as an author (Angela Paradis).

## Abstract

Motor neuron (MN) loss is a hallmark of neurodegenerative disorders, yet its assessment remains variable, confounding mechanistic and therapeutic interpretation. To address this, we conducted a systematic review and meta-analysis of spinal muscular atrophy (SMA) mouse studies, revealing 60% variability in reported MN loss, largely attributable to nonspecific spinal cord sampling. Using a whole-segment approach with tissue clearing, MN tracing, and multimodal imaging, we confirmed segment-dependent differences in MN counts. Common MN markers (SMI-32, Nissl) lacked specificity, whereas choline acetyltransferase (ChAT) provided robust labeling in murine and human spinal cords. Deep learning–based whole-mount segmentation enabled unbiased MN quantification and validated manual counts. Integrating analysis with computational modeling established segment sampling as a key driver of variability and revealed degeneration patterns: widespread MN loss in amyotrophic lateral sclerosis (ALS), selective MN loss in severe SMA, and preservation in mild SMA models. These findings establish a framework for reproducible MN quantification.

**Highlights:**

- Spinal cord segment-specific analysis reduces variability and allows accurate MN quantification
- ChAT is the most reliable MN marker in murine and human spinal cords
- Deep learning–based segmentation enables unbiased MN quantification in intact spinal cords
- MN degeneration is widespread in ALS but restricted to pools innervating proximal muscles in severe SMA

## Introduction

Spinal motor neurons (MNs) constitute the final common pathway of the motor system, driving essential functions including posture, locomotion, speech, and breathing. Their degeneration results in severe motor deficits, paralysis, and ultimately death ^1–3^. MN loss occurs physiologically during embryonic development as part of programmed cell death ^4,5^ but can also arise after trauma, ischemia, inflammation, as well as in neurodegenerative disorders such as poliomyelitis, hereditary motor neuropathies, and progressive muscular atrophy ^6–10^. MN degeneration is also a defining feature of MN diseases, including spinal muscular atrophy (SMA), amyotrophic lateral sclerosis (ALS), spinal and bulbar muscular atrophy (SBMA), and spinal muscular atrophy with respiratory distress type 1 (SMARD1) ^11–14^. Accurate and reproducible MN quantification is therefore essential for understanding the mechanisms governing MN survival and degeneration across development, injury, and disease.

Owing to the limited availability, variable quality and genetic heterogeneity of human postmortem tissue, mouse models are indispensable for longitudinal and mechanistic studies of MN degeneration. Several mouse models reliably recapitulate key features of disease and enable quantitative assessment of MN loss as a robust readout of disease severity, progression, and therapeutic response. However, the reported extent of MN loss varies widely even within the same disease mouse models ^15–20^, highlighting methodological variability and raising concerns about reproducibility, cross-study comparisons, and the interpretation of mechanistic and therapeutic outcomes.

A major source of variability may stem from inconsistent sampling of spinal cord regions containing disease-specific vulnerable versus resistant MN pools. The longitudinal organization of MNs into columns that innervate distinct muscle groups ^21–24^ underlies characteristic segmental vulnerabilities, with distal muscles preferentially affected in ALS and proximal muscles in SMA ^11,25–28^. Consequently, analyses targeting different spinal segments or MN pools may yield markedly different estimates of MN loss.

Beyond anatomical sampling, the molecular markers used to identify MNs represent another important source of variability in MN quantification. Classical Nissl and hematoxylin–eosin (H&E) staining have long been used to identify large ventral horn neurons presumed to be MNs but lack specificity, as they also label large non-MN neuronal populations within the ventral horn ^29–31^. SMI-32, an antibody recognizing non-phosphorylated neurofilament H, has also been utilized as an MN marker in rodents and humans, although accumulating evidence indicates expression in other ventral horn neuronal populations ^32–34^. More selective markers, including choline acetyltransferase (ChAT) and the homeobox protein HB9 are widely used but also label subsets of spinal interneurons outside MN pools ^35–37^, potentially confounding MN identification and the interpretation of experimental outcomes ^38,39^. Together, these limitations underscore the need for a systematic and quantitative comparison of commonly used MN markers to minimize errors in MN quantification.

Quantification strategies represent an additional source of variability in estimates of MN loss. While stereology-like methods aim to provide unbiased estimates of MN number, they require strict sampling parameters and are sensitive to tissue processing artifacts. Conversely, manual counting approaches are time-consuming, labor-intensive, and prone to inter-observer variability. Both approaches can be affected by developmental or disease-related changes in spinal cord size and morphology ^39^. Collectively, these challenges highlight the need to identify sources of variability and establish standardized approaches for MN quantification.

To address these issues, we performed a systematic literature review (SLR) and meta-analysis across SMA mouse models, identifying spinal cord segment selection as a major source of inter-study variability. We developed a standardized framework for reliable, reproducible, and unbiased MN quantification. To this end, we combined retrograde labeling with ChAT immunostaining, validated candidate MN-specific markers, assessed developmental and tissue-processing effects, and established both manual and machine-learning–based quantification approaches. Applying this framework revealed segment- and column-specific MN numbers in wild-type mice, as well as distinct selective MN vulnerabilities in ALS and SMA mouse models. Together, this study establishes a detailed, robust and scalable framework for standardized MN quantification across experimental paradigms and disease models.

## Results

### Meta-analysis identifies spinal segment sampling as a key driver for variability in reported motor neuron loss in SMA

Individual studies diverge markedly in the extent of reported MN loss, even when using identical SMA mouse models ^15–20^. To obtain an unbiased overview of this variability, we performed an SLR of SMA studies indexed in PubMed (**Table S1**). Using predefined population, intervention, comparator, outcomes, and study type (PICOS) criteria (**Table S2**), we identified 75 publications with data from the three most commonly used SMA mouse models: SMNΔ7, Smn^2B/–^, and Taiwanese (**Table S3**). The SMNΔ7 model was most frequently used (65% of studies), whereas Smn^2B/–^ and Taiwanese models accounted for ∼23% and 12% respectively (**Fig. S1A**). Across studies, the lumbar spinal cord was the most frequently examined region (82.5%), followed by the thoracic (11%) and cervical (6.5%) segments (**Fig. S1B**). ChAT (57%) immunostaining was the predominant method for MN quantification, with Nissl staining (20%), H&E (18%), HB9 (3%), SMI-32 and calcitonin gene-related peptide (CGRP) (1% each) being used less frequently (**Fig. S1C**). Remarkably, in these studies, reported MN loss varied widely (range: 0–60%) across all three models, and this variability persisted irrespective of the MN marker used or the spinal region analyzed (**Fig. 1A, B**). To determine whether this disparity in MN loss persists within a single spinal region, we further examined the lumbar spinal cord of SMNΔ7 mice, the most frequently studied experimental context. We subdivided lumbar MNs into pools innervating vulnerable proximal muscles (L1–L2 and the medial motor column at L5) and pools innervating more resistant distal muscles (L3–L6) ^40^. Studies that did not specify segmental levels were classified as general lumbar (**Fig. 1C**). This analysis revealed striking variability in reported MN loss (**Fig. 1C**), exposing potentially major inconsistencies in the SMA literature.

**Figure 1.**
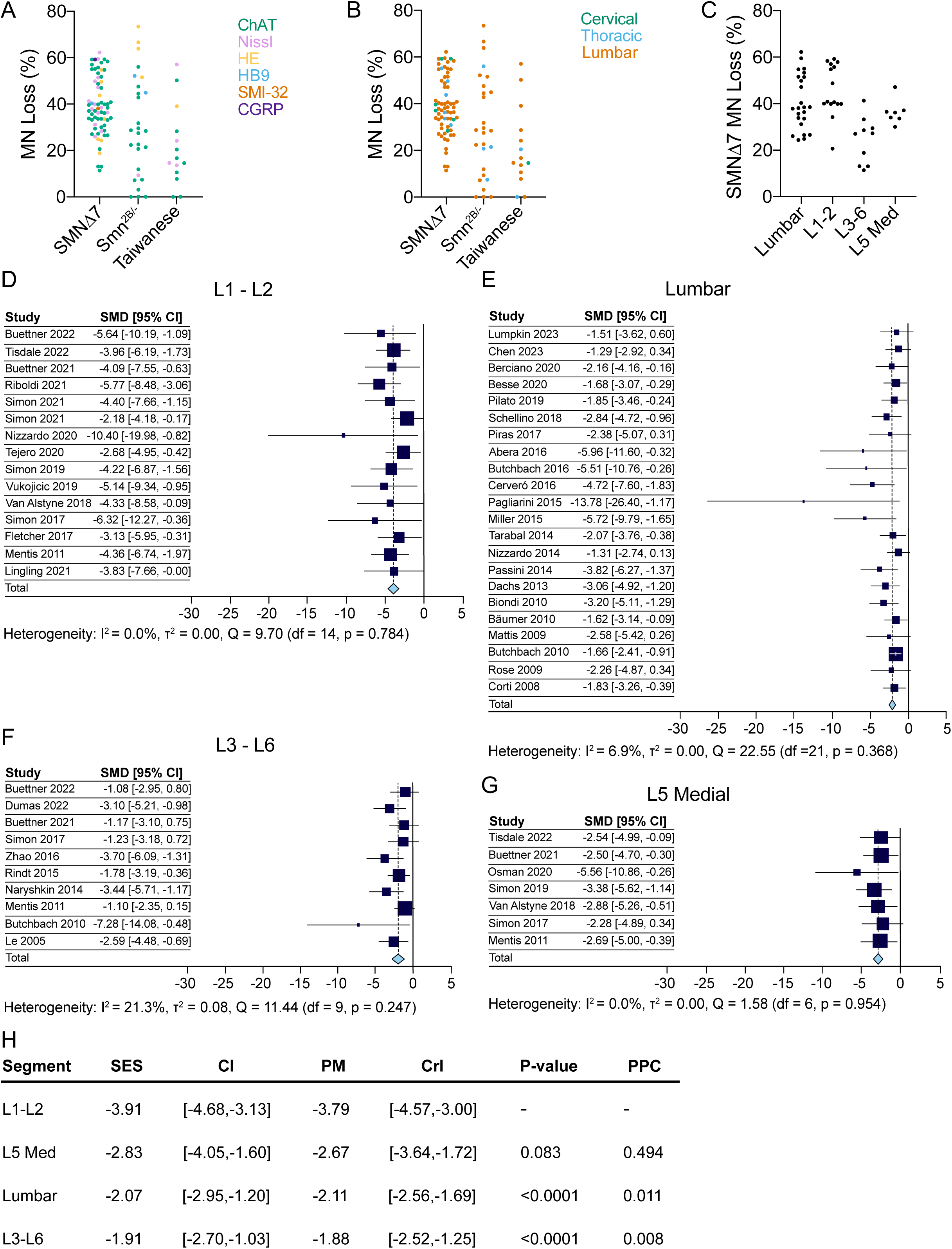
SLR and subgroup meta-analysis reveal spinal segment-dependent variability in MN loss in SMA mouse models. MN loss (%) across studies included in the SLR for three SMA models, SMNΔ7, Smn^2B/–^ and Taiwanese, grouped by MN marker (A) or spinal cord region analyzed (B). (C) MN loss (%) in the lumbar spinal cord of SMNΔ7 mice, subdivided into lumbar (segment not specified in the included studies), L1–L2, L3–L6 and L5 MMC. Summary of subgroup analyses for studies using the SMNΔ7 mouse model: L1–L2 (D), lumbar (E), L3–L6 (F) and L5 MMC (G). Blue boxes and black bars represent effect sizes and 95% confidence intervals, respectively. Larger boxes indicate greater study weight. Dotted lines indicate the summary effect size for each subgroup. SMD, standardized mean difference, a statistical measure equivalent to effect size; all subsequent references to effect size refer to this metric. I², estimated heterogeneity; τ², between-study variance in true effect size; Q (Cochran’s Q), heterogeneity statistic. (H) Comparison of subgroup-analysis estimates with a Bayesian model. SES, summary effect size; CI, 95% confidence interval; PM, posterior mean; CrI, credible interval; PC, pairwise comparison; PPC, posterior probability of contrast.

We next wanted to address the nature of the marked variability in MN loss observed in the SLR by performing a meta-analysis restricted to the lumbar spinal cord of the SMNΔ7 mouse model, integrating 43 publications comprising 54 lumbar region data groupings. The effect sizes derived from control and SMA MN counts showed low overall heterogeneity (I² = 16.2%, Q(df=53) = 66.0499, p = 0.1075), though the bootstrapped upper 95% confidence interval bound approached 51.5%, suggesting moderate heterogeneity. We next considered which factors could be acting as confounders hindering study comparability. Immunohistochemical staining and quantification methods presented a significant source of variability between studies, but aspects regarding staining quality and marker combinations prevented further consideration in this analysis. Another factor that varied among the reporting was the spinal segment. This was controlled for in the subsequent subgroup analysis by spinal segment (Lumbar, L1–L2, L3–L6, L5 medial motor columns (MMC)), which revealed a significantly improved model fit (Q(3) = 19.55, p = 0.0002; **S1F**) and markedly reduced intra-study heterogeneity (**Fig. 1D-G**). The test of moderators of this model was significant (I² = 0.0%, Q(M)(df=3) = 20.7736, p=0.0001), indicating that the explanation of heterogeneity was likely due to the spinal region. Segment-specific analyses confirmed minimal overlap of the effect size confidence intervals across groups (**Fig. 1H**), indicating a unique relation between MN loss and spinal segment. Furthermore, Bayesian multilevel modeling with segment identity as a control reinforced these findings by revealing comparatively narrowed credible intervals (**Fig. S1E**). Additional analyses with pairwise comparisons and Bayesian posterior probabilities showed that “L1–L2” contained the strongest relation to MN loss, clearly distinct from either “Lumbar” or “L3–L6 segments”, whereas L5 MMC exhibited a weaker but consistent trend (**Fig. 1H, S1D-E**).

Together, meta-analytic and Bayesian analyses identify L1–L2 MN pools as the most vulnerable and reveal segmental sampling as a key driver of variability in reported MN loss across SMA studies.

### Divergent motor neuron distribution across the spinal cord revealed by segment-specific analysis

To translate our meta-analysis findings into experimental validation, we first sought to establish a reliable, segment-specific method for quantifying MNs in spinal cord tissue. To map segmental variation in MN numbers, we dissected spinal cords from four-day-old (P4) C57BL/6 mice and retrogradely filled L1–L6 ventral roots *ex vivo* with the fluorescent dyes Fluorescein Dextran and Texas Red in an alternating manner (**Fig. 2A** and **Fig. S2A, C, D**). Spinal cords were subsequently immersion-fixed with 4% paraformaldehyde (PFA), ChAT-immunostained, cleared (**Fig. S2B**), and imaged in whole-mount (the intact spinal cord without any sectioning) by confocal microscopy (Approach 1, **Fig. 2A, B**). To directly compare MN quantification in intact spinal cords and sections, we washed these cleared lumbar spinal cords in PBS after the first imaging session to reverse clearing, sectioned them into 75 µm-thick transverse slices using a vibratome, and subsequently re-imaged them (Approach 2, **Fig. 2A, B**). Retrograde ex vivo labeling in whole-mount and sectioned preparations revealed minimal rostro–caudal overlap between adjacent MN pools, restricted to single sections, enabling precise assignment of MNs to specific segments (**Fig. 2B** and **Fig. S2D**). While 60% of ChAT⁺ MNs were retrogradely filled, nearly all (98%) labeled MNs were ChAT⁺, validating ChAT immunoreactivity as a reliable marker of MN identity (**Fig. S2E–G**). Along the span of L2, MNs segregated into lateral motor columns (LMC) and MMC, with LMC numbers increasing and MMC numbers decreasing along the rostro-caudal axis with the exception of L6, in which MNs formed small, discrete pools rather than large continuous columns (**Fig. 2B, C**). Manual segmental analysis showed MN numbers increasing from ∼300 in the rostral lumbar segment L1 to ∼700 the in caudal lumbar segment L5 before declining to ∼300 in L6 in both approaches (**Fig. 2B–D**). These consistent MN counts in whole-mount and sectioned cords validate the retrograde spinal root-labeling with ChAT as a reliable quantification method and demonstrate that lumbar sections cannot be pooled together due to segment-specific differences.

**Figure 2.**
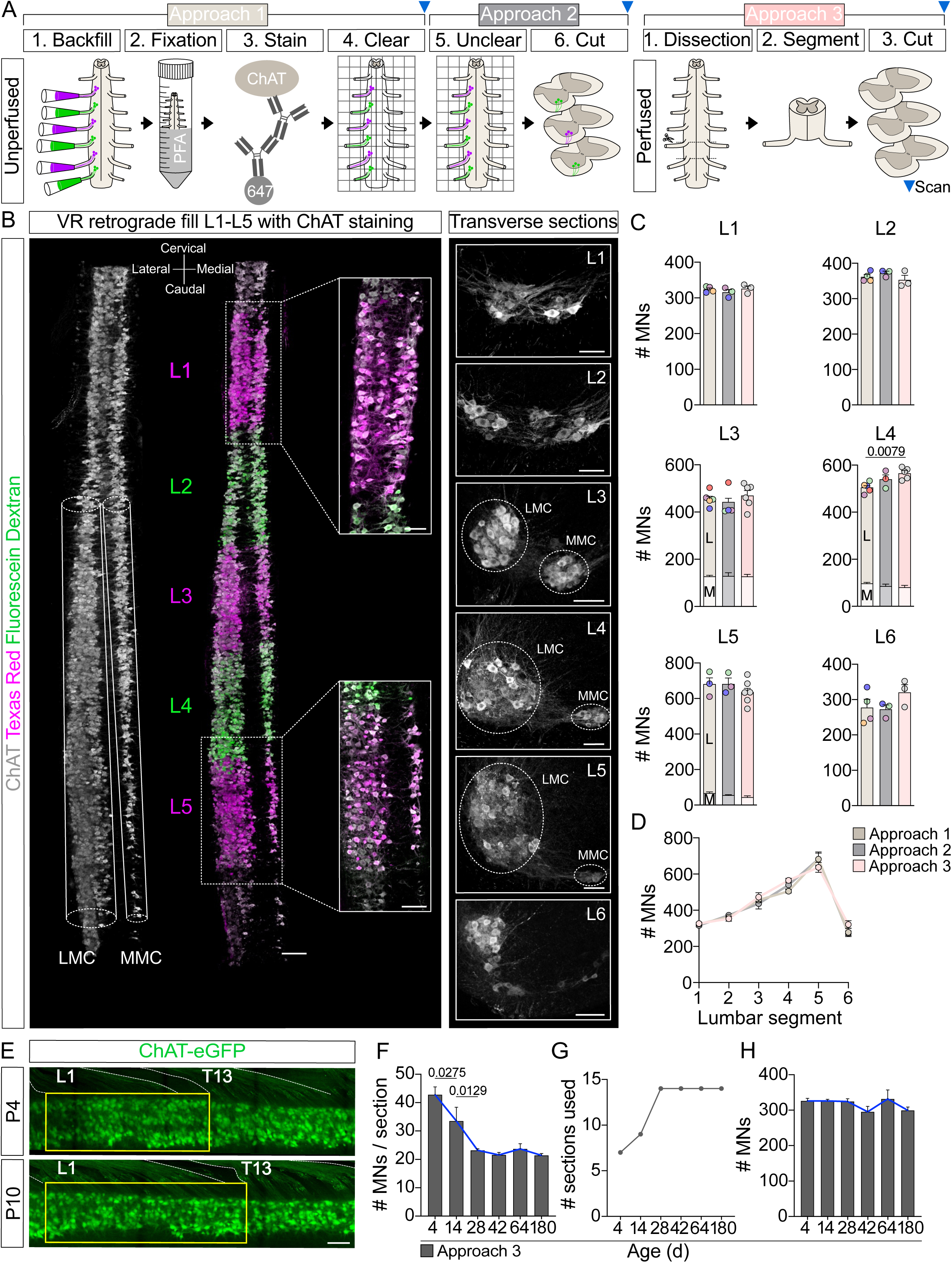
Sectioning-based workflow enables reproducible, segment-specific quantification of lumbar MNs. (A) Schematic of the clearing workflow (Approaches 1 and 2) and dissection workflow (Approach 3). Blue arrows indicate the imaging time point within each approach. (B) Confocal image of a ChAT-stained (gray), cleared intact lumbar spinal cord from a P4 C57BL/6 mouse with retrogradely labeled segments in alternating order of Texas Red (magenta) and Fluorescein Dextran (green). Scale bar = 200 µm. Highlighted insets show higher magnification views of L1 and L5 segments. Transverse sections of ChAT-stained spinal cord show individual lumbar segments, with the MMC and LMC indicated. Scale bars = 50 µm. (C) Quantification of MNs in L1–L6 of the intact lumbar spinal cord (Approach 1), vibratome-sectioned segments of the same spinal cord identified by retrograde labeling (Approach 2), and sections of individual spinal cord segments identified by ventral roots (Approach 3). Colored points represent individual animals, with matching colors indicating the same animal across Approaches 1 and 2. For each segment, n values are listed in the order of Approach 1, Approach 2, and Approach 3: L1: n = 4, 3, 3; L2: n = 4, 3, 3; L3: n = 5, 4, 6; L4: n = 4, 3, 5; L5: n = 3, 3, 6; L6: n = 4, 3, 3. L (LMC) and M (MMC) denote MN counts from the respective motor columns. (D) Quantification of MNs from the same groups as in (C), plotted across lumbar spinal segments. For segments L3–L5, LMC and MMC counts were combined. (E) T13 to L1 MN pool from a cleared whole-mount spinal cord of P4 and P10 ChAT-eGFP mice. T13 and L1 ventral roots are outlined in white. The yellow box indicates the L1 spinal cord segment as defined by ventral root position. Scale bar = 100 µm. (F) Number of MNs per vibratome section (75 µm), (G) number of vibratome sections spanning the entire L1 segment, and (H) total number of MNs in the L1 segment from C57BL/6 mice at P4, P14, P28, P42, and P180, as identified by ventral root dissection (Approach 3). P4 and P180 data points are from Figure 1C and S6K. (n = 3 unless otherwise indicated; P180, n = 4). Statistical analysis was performed using one-way ANOVA (C: L1, L2, L4, L5; F, H) and Kruskal–Wallis test (C: L3) comparing the total number of MNs.

To validate a less labor-intensive approach for segment-specific quantification without retrograde labeling ^20,26,41,42^, we perfused P4 C57BL/6 mice and dissected lumbar segments using ventral root landmarks as described previously ^43^ (**Fig. S2A**). Each segment was sectioned in its entirety at 75 µm thickness with a vibratome, stained for ChAT, and analyzed for MNs by confocal microscopy (Approach 3, **Fig. 2A**). MN numbers of each segment were found to be indistinguishable between Approaches 1 and 2, establishing this sectional method as a practical and reproducible tool for routine, segment-specific MN quantification (**Fig. 2C, D**).

To evaluate the reliability of this method throughout the spinal cord development and to determine the impact of maturation upon MN column dimensions, we cleared perfused whole spinal cords from P4 and P10 mice, which expressed eGFP under the ChAT promoter (ChAT-eGFP). Confocal scans of the whole-mount T13–L2 region suggested that total MN numbers remained stable between P4 and P10, but, as the L1 segment lengthened during postnatal development, MN density decreased (**Fig. 2E**). To quantify these differences, we compared MN numbers per section, MNs per segment, and sections per segment in ChAT-stained and retrogradely labeled L1 segments of FVB control mice of the SMNΔ7 mouse line (**Fig. S2I-K**). While the number of MNs per section decreased from P4 to P13 (**Fig. S2I**), the length of the developing spinal segment increased, as reflected by a higher number of sections per segment (**Fig. S2J**), resulting in stable MN numbers across ages (**Fig. S2K**). A similar – though slightly delayed – trend was observed in C57BL/6 mice when L1 segments were delineated using ventral root landmarks. Here, MN numbers per section decreased from P4 through P14 to P28, accompanied by increasing segmental length, but reached a plateau at later time points (**Fig. 2F-H**), indicating completion of the maturation process after the fourth postnatal week. Importantly, the total number of MNs per L1 segment was consistent across ages (**Fig. 2H** and **Fig. S2K**), demonstrating that the quantification of all MNs within an entire spinal segment provides a reliable metric across developmental stages.

Overall, whole-mount spinal cord analysis reveals a rostro-caudal gradient in lumbar MN numbers and establishes a reliable ChAT-based method for segment-specific, manual quantification.

### Spinal interneurons undermine the specificity of SMI-32 and Nissl as motor neuron markers

After validating the reliability of ChAT immunostaining in marking MNs, we assessed the specificity of other commonly used markers (**Fig. 1A** and **Fig. S1C**), using ventral roots as landmarks as defined by Approach 3 (**Fig. 2A**). We performed quadruple labeling with antibodies against ChAT, HB9, SMI-32, and Nissl on spinal cords from P42 C57BL/6 mice focusing on L1 and L5 segments that innervate proximal/axial and distal muscles, respectively^40^. HB9 and Nissl each labeled ∼90-95% of all ChAT⁺ MNs, while SMI-32 labeled ∼80% in both segments (**Fig. 3A, B** and **Fig. S3A, B**). Importantly, while HB9 selectively marked both α- and γ-MNs (**Fig. S3E**), SMI-32 and Nissl exhibited substantial off-target labeling in the ventral horn, detecting ∼160% (SMI-32) and ∼280% (Nissl) additional non-MNs – i.e. spinal interneurons – in L1 and ∼75% and ∼120% in L5 in mice, respectively (**Fig. 3A, C** and **Fig. S3A, C, D**). Extending these findings from mice to human autopsy tissue, we analyzed 20 µm cryosections of postmortem thoracic spinal cord tissue. Nissl and SMI-32 labeling identified ∼70–80% of ChAT⁺ MNs, while both markers exhibited substantially fewer off-target profiles in humans (∼20–40%) compared to mice (**Fig. 3D–F**).

**Figure 3.**
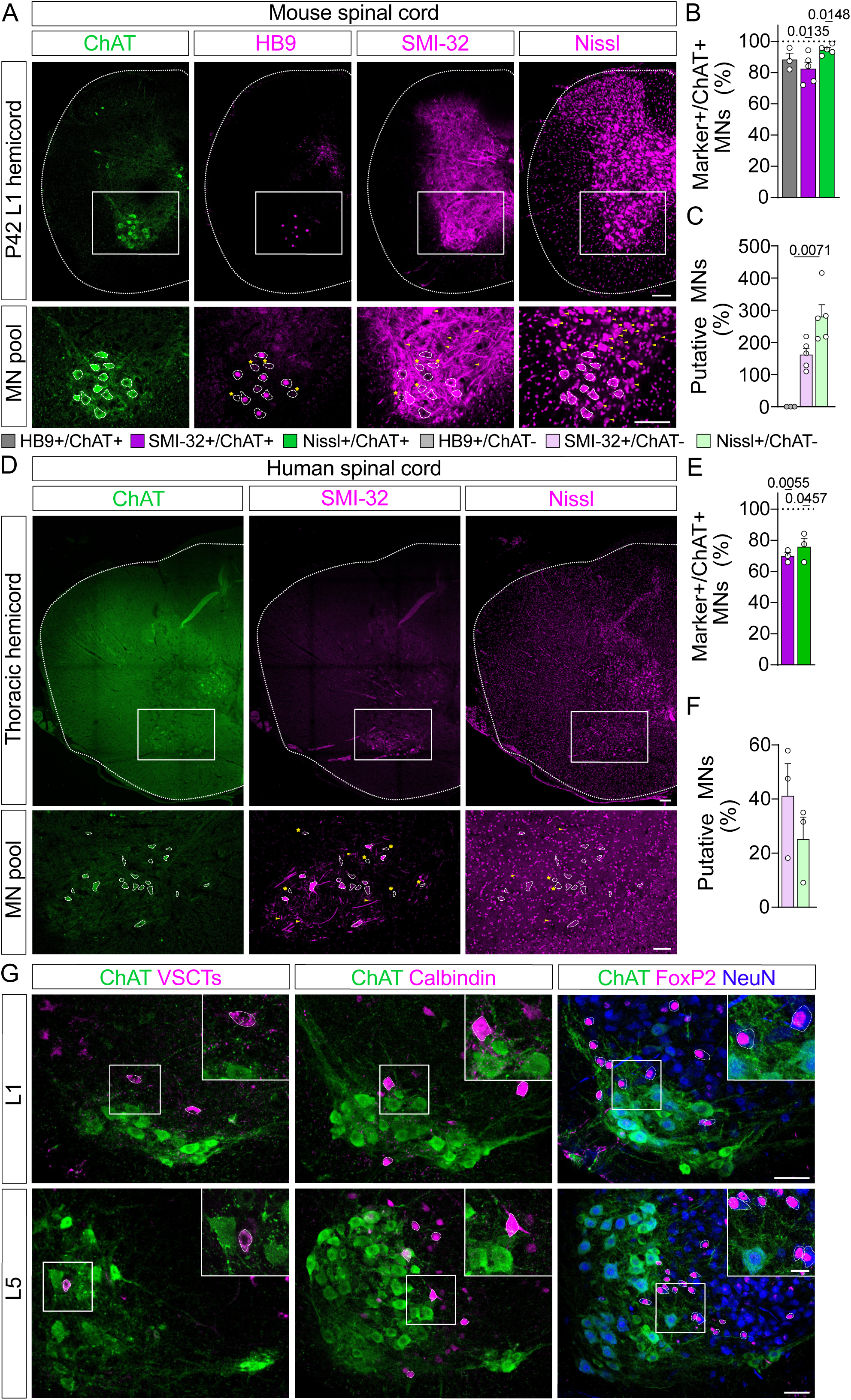
Marker non-specificity and dense interneuron proximity lead to inaccurate MN labeling. (A) Immunostaining of L1 hemicord (upper row) and MN pool (lower row) for ChAT (green), SMI-32, Nissl, and HB9 (magenta) from a P42 C57BL/6 mouse. Scale bars = 100 µm. Yellow asterisks indicate marker-ChAT+ MNs, and yellow arrows indicate marker+ ChAT-putative MNs. MN shapes, as identified by ChAT, are shown as white dotted outlines. (B, C) Quantification of SMI-32, Nissl, and HB9 labeling in ChAT+ MNs (B) and in ChAT-cells classified as putative MNs (C), expressed as percentages within the L1 ventral horn surrounding the MN pool of P42 C57BL/6 mice (n = 3 for HB9; n = 5 for SMI-32 and Nissl), as indicated by the white box in (A). (D) Immunostaining of thoracic hemicord (upper row) and MN pool (lower row) for ChAT (green), SMI-32, and Nissl (magenta) from a 9-month-old human spinal cord sample. Scale bars = 200 µm (upper) and 100 µm (lower). Yellow asterisks indicate marker-ChAT+ MNs, and yellow arrows indicate marker+ ChAT-putative MNs. MN outlines identified by ChAT are shown as white dotted outlines. (E, F) Quantification of SMI-32 and Nissl labeling in human thoracic spinal cord samples in ChAT+ MNs (E) and in ChAT-cells (putative MNs) (F), expressed as percentages within the ventral horn surrounding the MN pool (n = 3), as indicated by the white box in (D). (G) Interneurons (magenta) in the ventral horn identified by CTB injection into the cerebellar vermis to label VSCTs, Calbindin for Renshaw cells, and FoxP2 for V1 interneurons, co-stained with ChAT (green) in L1 (upper row) and L5 (lower row) of P4 C57BL/6 mice. NeuN (blue) is co-labeled in the FoxP2 row to visualize neuronal somata. Insets show higher magnification of interneurons in close proximity to MNs. Scale bars = 50 µm and 20 µm for insets. Outlines of interneurons are shown as white dotted lines. Statistical analysis was performed using one-sample t-tests against ChAT+ MN values (B, E), Kruskal–Wallis test (C), or unpaired t-test (F).

To identify large ChAT-neurons located within or adjacent to MN pools, we immunolabeled subsets of ventral interneurons, including ventral spinocerebellar tract (VSCT) neurons identified by injecting cholera toxin B subunit (CTB+) in the cerebellum, calbindin+ Renshaw cells, and FoxP2+ V1 interneurons in L1 and L5 spinal segments of P4 C57BL/6 mice ^31,44,45^. Some interneurons were similar in size to small MNs, whereas others, such as VSCT neurons, were MN–sized and located adjacent to or within MN pools (**Fig. 3G**).

Next, we addressed whether different developmental stages and tissue fixation during processing could introduce variability in MN counts. To assess potential fixation effects on MN counts during maturation, we examined spinal cords from P10 and P60 ChAT-eGFP mice, rapidly dissected in ∼4 °C oxygenated aCSF with subsequent immersion-fixation with 4% PFA. L1 segments, identified by ventral root landmarks, were sectioned at a vibratome. At P10, GFP-expressing MNs in immersion-fixed tissue revealed nearly complete colocalization with ChAT and HB9 and partial colocalization with SMI-32 (**Fig. S3F**). In contrast, immersion-fixed P60 spinal cords lacked detectable GFP, even with antibody enhancement or ChAT signals in MNs, whereas HB9 and SMI-32 remained visible (**Fig. S3F**). By comparison, transcardially perfused adult spinal cords exhibited robust ChAT staining (**Fig. S3A-D**), indicating that cytoplasmic ChAT degrades more rapidly than nuclear HB9 or cytoskeleton-enriched SMI-32 in immersion-fixed adult spinal cords, underscoring the necessity of perfusion for MN-specific labeling with ChAT in adult, fully myelinated tissue.

These findings identify specific interneurons as a likely source of mistakenly assigned MNs when using nonspecific markers such as SMI-32 or Nissl, highlight the importance of perfusion, and demonstrate HB9 and ChAT as highly specific markers for accurate, manual MN quantification.

### Deep learning–based segmentation enables unbiased motor neuron quantification within the intact spinal cord

In addition to tissue fixation and marker selection, variability in sectioning methods and manual MN quantification across investigators constitute potential sources of error. To evaluate these possibilities, we first examined how vibratome versus cryostat sectioning affects MN integrity and counts. We sectioned entire L1 spinal segments from perfused P10 FVB control mice of the SMNΔ7 line using both methods, followed by ChAT immunofluorescence. As expected, 75 µm vibratome sections consistently contained higher MN counts than 20 µm cryostat sections (**Fig. 4A, B**). To correct for overestimation due to nuclei split across adjacent sections, we measured the MN nuclear diameter (**Fig. S4A**) and applied the Abercrombie correction ^46^. MN counts in thicker vibratome sections were reduced by only ∼10%, whereas thin cryostat sections required ∼35% adjustment (**Fig. 4C, D**). To test whether this variability persists under neurodegenerative conditions, we performed the same analysis in end-stage P10 SMNΔ7 mutant littermates ^18^. Both methods reliably detected a ∼50% reduction in L1 MNs per section in SMNΔ7 mice compared to controls, consistent with previous reports ^26,42,47–50^. This loss is independent of the Abercrombie correction, as MN nuclear diameters are unaltered in SMA (**Fig. 4A–E** and **Fig. S4A**). These findings indicate that both sectioning methods detect MN loss similarly; however, thicker vibratome sections are less affected by double-counting and may provide MN counts that more closely reflect those of the intact spinal cord.

**Figure 4.**
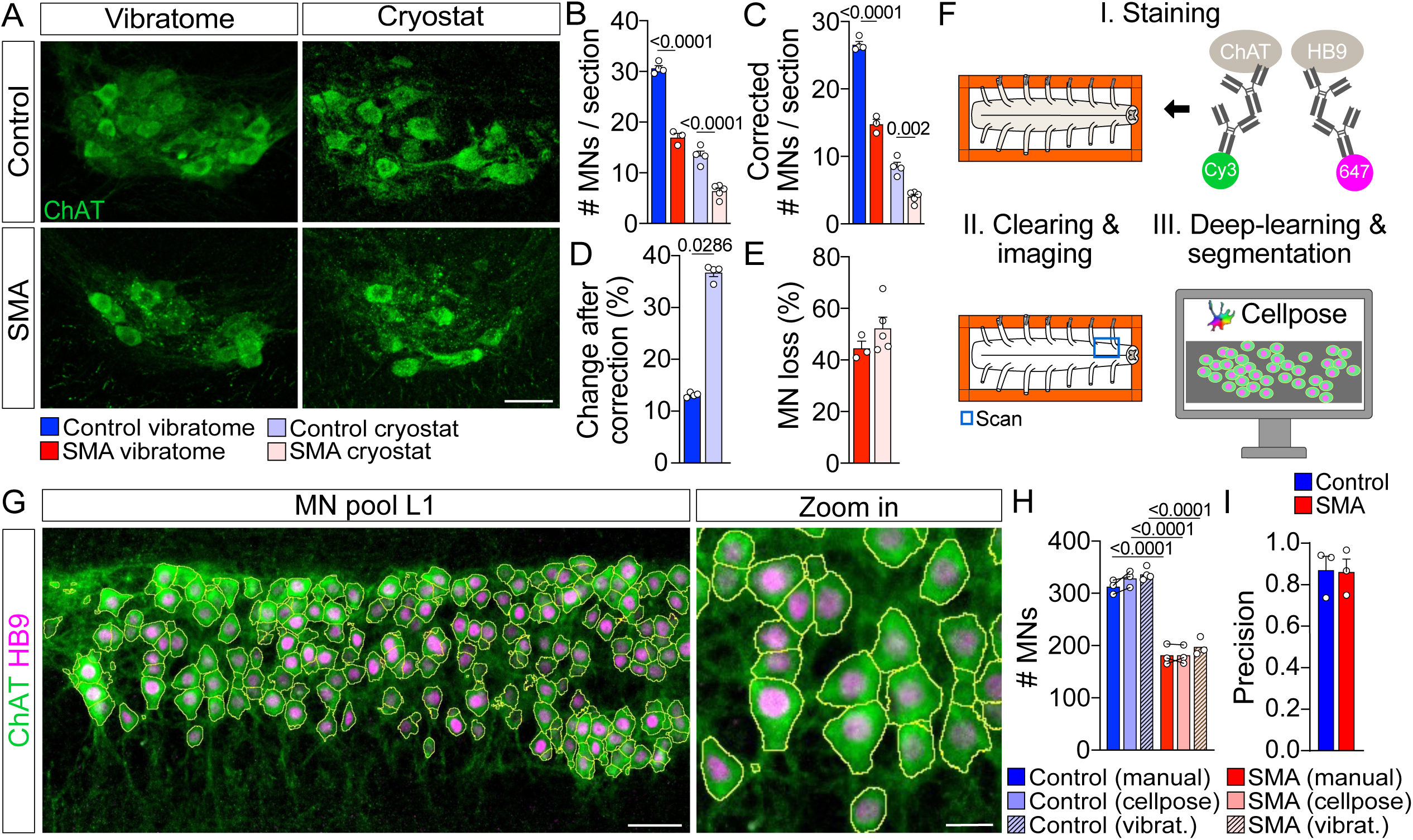
Deep learning–based Cellpose automated MN quantification confirms manual vibratome-based counts. (A) Immunostaining of the L1 MN pool for ChAT (green) in sections cut using a vibratome (75 µm) or cryostat (20 µm) from end-stage control and SMNΔ7 mice. Scale bar = 50 µm. (B) Quantification of MNs per section for the groups shown in (A) (for both sectioning methods, control: n = 4, SMA vibratome: n = 3, SMA cryostat: n = 5). (C) Quantification of Abercrombie-corrected MNs per section for the same groups as in (B). (D) Percent change after Abercrombie correction for both cryostat and vibratome control groups. (E) Calculated MN loss (%) between SMA and control animals for the same groups as in (B) (vibratome: n = 3; cryostat: n = 5). (F) Schematic of the workflow for automated analysis using Cellpose of whole-mount, immunostained, cleared spinal cords. The blue box indicates the scanned area of the L1 segment. (G) Immunostaining of cleared whole-mount L1 MN pool for ChAT (green) and HB9 (magenta) from a C57BL/6 mouse at P4, segmented with Cellpose (yellow outlines). Scale bars = 50 µm and 20 µm. (H) Quantification of MNs in the entire L1 segment of P10 control and SMNΔ7 mice (n = 3) using manual and Cellpose-based segmentation, as well as from L1 segments cut using a vibratome (control: n = 4; SMA: n = 3). Connecting lines indicate measurements from the same animal for manual and Cellpose-based quantification. (I) Calculated precision of Cellpose MN quantification in the L1 segment of P10 control and SMNΔ7 mice (n = 3), compared to ground-truth manual quantification. TP = true positives, FP = false positives, FN = false negatives. Statistical analysis was performed using one-way ANOVA (B, C, H), Mann–Whitney test (D), and unpaired t-test (E, I).

To directly compare vibratome section-based manual counts with the true MN population of the intact spinal cord, we applied a deep-learning, automated pipeline for unbiased, whole-tissue quantification (**Fig. 4F**). To do so, we dissected spinal cords of perfused P10 FVB control and SMNΔ7 mice and pinned them into a 3D printer custom-made chamber (**Fig. S4B**). Subsequently, we immunolabeled the dissected spinal cord for ChAT and HB9, cleared the tissue, and imaged the entire L1 segment identified using ventral root landmarks. The confocal stacks were uploaded into the open-source, deep-learning software Cellpose ^51^. Representative images of these stacks were used to train an existing Cellpose model ^52^ to segment and quantify MNs based on ChAT signal with additional help of HB9 nuclear labeling (**Fig. 4F, G** and **Supp. Fig. 4C**). The trained Cellpose model generated MN counts that closely matched manual assessments of the same stacks and parallel vibratome L1 sections without Abercrombie correction, accurately reflecting the degree of MN loss (**Fig. 4H**). Quantification of MNs with Cellpose achieved a precision of 0.86, indicating that the majority of automatically detected MNs corresponded to manually identified MNs with few false-positive detections in both control and SMA mice (**Fig. 4I**). This automated approach establishes an unbiased method for reliable MN quantification of intact spinal cord tissue and validates the manual vibratome sectional analysis under both healthy and disease conditions.

### Random sampling masks column-specific variability of MN loss across spinal columns in SMA

Our meta-analysis identified inconsistent segment sampling as a key variable in reported MN loss across SMA studies (**Fig. 1**). With a validated ChAT-based sectional approach in place, we next examined the consequences of random sampling by quantifying MN loss across the entire lumbar spinal cord of control and end-stage SMNΔ7 mice by integrating these data into a computational model (**Fig. 5A**). We applied ChAT and NeuN immunolabeling on 75 µm thick L1 to L6 vibratome sections to distinguish α-MNs innervating extrafusal muscle fibers (ChAT+, NeuN+) from γ-MNs innervating intrafusal fibers (ChAT+, NeuN-) ^53,54^. Whereas L1–L2 and MMC MNs across L3–L5 innervate proximal and axial muscles, LMC MNs and dispersed L6 pools innervate distal hindlimb muscles ^40^. γ-MNs were spared in all distal muscle–innervating pools with trends for modest reductions observed in proximal muscle–innervating pools. Degeneration remained restricted to α-MNs innervating proximal and axial muscles throughout the lumbar spinal cord with severity decreasing from upper to lower lumbar segments (**Fig. 5B–H**).

**Figure 5.**
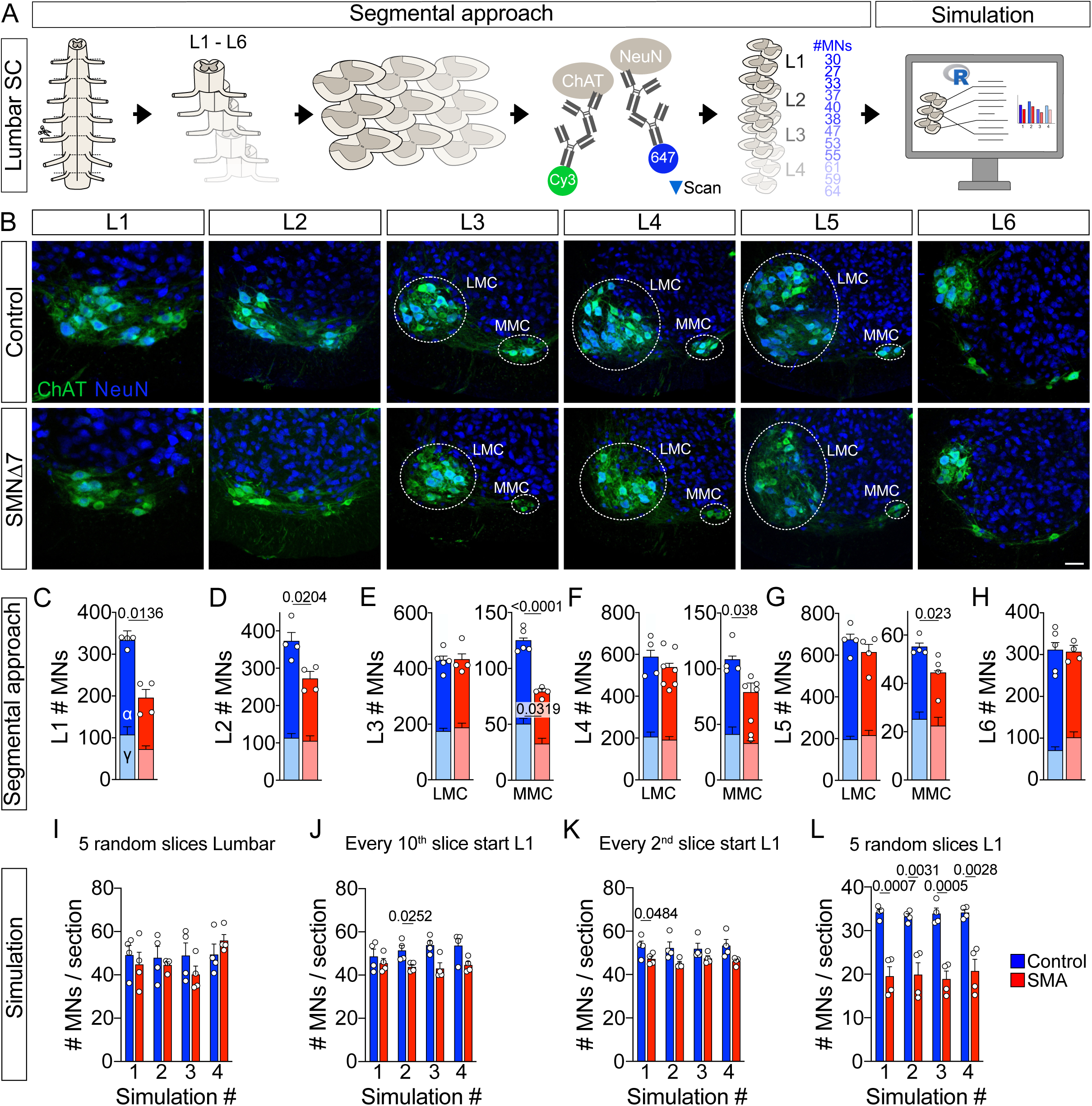
Non-segmented counting approaches mask selective MN vulnerability in severe SMA mice. (A) Schematic of the workflow for manual segment-specific MN quantification in lumbar spinal cords, followed by simulation of randomized segmental sectioning implemented in R. (B) Immunostaining of L1–L6 spinal segments for ChAT (green) and NeuN (blue) from P11 control (upper row) and SMNΔ7 (lower row) mice. ChAT+/NeuN− = γ-MNs; ChAT+/NeuN+ = α-MNs. Scale bar = 50 µm. LMC and MMC are outlined in L3–L5 with white dotted lines. (C–H) Quantification of MN numbers in L1 (C), L2 (D), L3 (E), L4 (F), L5 (G), and L6 (H) spinal segments of P11 control and SMNΔ7 mice (L1: n = 4 per group; L2: n = 4 per group; L3: control n = 5, SMNΔ7 n = 4; L4: control n = 4, SMNΔ7 n = 7; L5: n = 4 per group; L6: control n = 6, SMNΔ7 n = 4). Segments L3–L5 are further subdivided into LMC and MMC. (I–L) Results of randomization simulations of MN number per slice in control versus SMA mice under different sampling conditions: (I) five random slices from L1–L6 (lumbar); (J) every 10th slice after a random start within L1; (K) every 2nd slice after a random start within L1; and (L) five random slices from L1 (n = 4 per group). Four simulation rounds are shown (1–4). Statistical analysis was performed using unpaired t-test (C–H, I–L), Welch’s t-test (F, MMC α-MNs), and Mann–Whitney test (H α-MNs; I, round 2; J, rounds 3–4; K, rounds 2–3).

Next, we incorporated these newly acquired ChAT+ MN counts across the entire lumbar spinal cord into an R-based modeling script (**Fig. 5A** and **I**-L, Fig. S5) to simulate random stereology-like sampling of MN quantification, as performed in previous studies ^55–57^. Four rounds of simulated random sampling of five consecutive lumbar sections failed to detect significant MN loss between control and SMNΔ7 animals (**Fig. 5I**). Similarly, sampling every 2nd or 10th section throughout the lumbar spinal cord, starting at a random point in L1, produced a significant difference in only one instance, with a minor effect size (**Fig. 5J, K**). In contrast, sampling five random sections from L1 or L2 consistently detected significant MN loss in all four simulation rounds (**Fig. 5L** and **Fig. S5**).

These findings highlight that random sampling across lumbar spinal cord segments carries a high risk of misestimating MN loss in SMA due to segment- and pool-specific heterogeneity in degeneration.

### Column-specific MN vulnerability differs among mouse models of MN disease

Our meta-analysis (**Fig. 1**) and random sampling (**Fig. 5**) suggested that the selective vulnerability of MN pools innervating proximal/axial muscles accounts for lumbar MN loss in severe SMNΔ7 mice. To determine whether this pattern extends beyond the lumbar cord, we applied ChAT and NeuN immunostaining to perfused end-stage P10 SMNΔ7 mice to quantify MN loss in cervical (C6) and thoracic (T9) segments which primarily innervate proximal forelimb and abdominal/trunk muscles, respectively (**Fig. 6A**). In accord with our observation within the lumbar spinal cord, C6 and T9 MN pools exhibited a significant (∼30%) α-MN loss and an additional ∼30% reduction in γ-MNs in T9 (**Fig. 6A–C**). Consistent with a previous report in the same mouse model ^26^, neither γ-MNs nor α-MNs exhibited significantly reduced soma size in SMNΔ7 mice (**Fig. S6A-D)**. In contrast to these observations in severe SMA mice, human autopsy spinal cords from SMA type I patients showed a ∼50% reduction in α-MN soma size, whereas γ-MNs showed a ∼25% reduction that did not reach statistical significance (**Fig. S6E-H**). Together, these data reveal a consistent, strong loss of α-MNs innervating proximal and axial muscles and modest vulnerability of γ-MNs at distinct spinal cord levels in severe SMNΔ7 mice.

**Figure 6.**
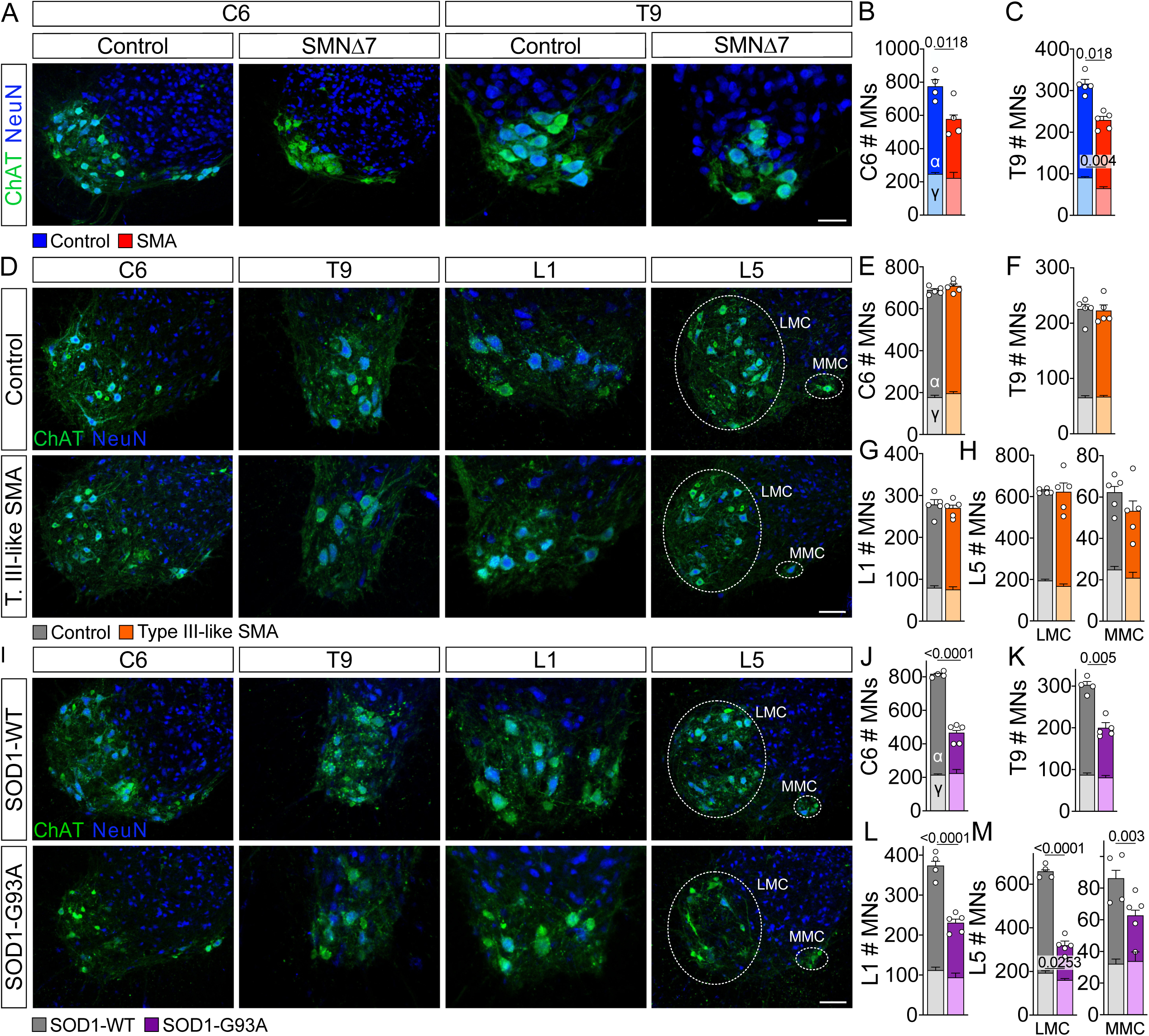
Differential vulnerability of MN pools across mouse models of motor neuron disease. (A) Immunostaining of C6 and T9 spinal segments for ChAT (green) and NeuN (blue) in P11 control (upper row) and SMNΔ7 (lower row) mice. ChAT+/NeuN− cells correspond to γ-MNs, whereas ChAT+/NeuN+ cells correspond to α-MNs. Scale bar = 100 µm in C6 and 40 µm in T9. (B, C) Quantification of MN pools in C6 (B) and T9 (C) segments of P11 control and SMNΔ7 mice (C6: n = 4 per group; T9: n = 5 per group). (D) Immunostaining of C6, T9, L1, and L5 spinal segments for ChAT (green) and NeuN (blue) in P270 control (upper row) and Type III-like SMA (lower row) mice. Scale bar = 100 µm in C6 and L5, and 50 µm in L1 and T9. In L5, the LMC and MMC are outlined with white dotted lines. (E–H) Quantification of MN pools in C6 (E), T9 (F), L1 (G), and L5 LMC and MMC (H) segments of P270 control and Type III-like SMA mice (n = 5 per group). (I) Immunostaining of C6, T9, L1, and L5 spinal segments for ChAT (green) and NeuN (blue) in P130 SOD1-WT (upper row) and SOD1-G93A (lower row) mice. Scale bar = 100 µm in C6 and L5, and 50 µm in L1 and T9. In L5, the LMC and MMC are outlined with white dotted lines. (J–M) Quantification of MN pools in C6 (J), T9 (K), L1 (L), and L5 LMC and MMC (M) segments of P130 SOD1-WT and SOD1-G93A mice (control: n = 4; ALS: n = 5). Statistical analysis was performed using an unpaired t-test (B, C, E–H, J–M), a Mann–Whitney test (F α-MNs, H LMC γ-MNs, K γ-MNs), or Welch’s t-test (G γ-MNs, H LMC α-MNs).

Next, we tested whether the selective vulnerability of α-MNs innervating proximal muscles is conserved in an adult-onset, 4-copy SMN2 Type III-like SMA mouse model, which has been reported to exhibit ∼30% lumbar MN loss at P270 on a C57BL/6 background based on analyses with non-specific MN markers ^58^. To investigate this, we analyzed both α- and γ-MNs using ChAT and NeuN in C6, T9, L1, and L5 segments isolated from perfused Type III-like SMA mutant mice and age-matched controls at P180 and P270. Strikingly, we found that α- and γ-MN numbers were fully preserved across all segments and ages (**Fig. 6D-H and Fig. S6I-L**). Consistent with this, p53 pathway activation, which drives MN death in severe and intermediate SMA mouse models ^20,42,48,59^, was completely absent in the nuclei of L1 MNs of Type III-like SMA mice (**Fig. S6M**). Accordingly, neuromuscular junctions (NMJs) in the distal tibialis anterior and axial quadratus lumborum muscles were fully innervated (**Fig. S6N–P**), unlike in severe and intermediate SMA mouse models ^20^. Together, these findings demonstrate the absence of MN loss in the 4-copy SMN2 Type III-like mouse model of SMA.

Finally, we investigated segment-specific MN loss in another adult-onset MN disease, amyotrophic lateral sclerosis (ALS), using the established SOD1-G93A mouse model ^28^. We analyzed MNs in the C6, T9, L1, and L5 segments at end-stage (P130) in SOD1-G93A mutants and SOD1-WT controls. In SOD1-G93A mutants, α-MNs exhibited substantial degeneration (>50%) across all segments, with the distal muscle–innervating L5 LMC MNs showing the greatest loss (**Fig. 6I-M**), consistent with the distal-onset phenotype characteristic of ALS rodent models ^11,27,28^. We also found that α-MNs were significantly reduced in size in this mouse model (**Fig. S6Q-S**). In contrast, γ-MNs remained largely unaffected in numbers and size (**Fig. S6T**), in accordance with previous reports ^60,61^.

Taken together, segment-specific analyses highlight the importance of specific markers in revealing extensive α-MN loss in SOD1-G93A mice, selective proximal vulnerability in severe SMNΔ7 mice, and the absence of MN degeneration in the Type III–like SMA model.

## Discussion

MN degeneration is a hallmark of several neurodegenerative disorders, yet reproducible quantification of MN loss remains technically challenging, limiting accurate disease phenotyping and assessment of therapeutic efficacy in preclinical animal studies. We identified substantial variability in reported MN loss across studies using identical SMA mouse models, primarily due to inconsistent spinal cord segment sampling with additional contributions from tissue processing and MN marker selection. To overcome these limitations, we demonstrated the superior accuracy of a segment-specific, ChAT-based manual sectioning strategy, validated using unbiased, automated machine learning–based whole-mount counting. Applied to SMA and ALS mouse models, this approach revealed disease-specific spatial selective MN vulnerability, demonstrating that methodological rather than biological differences underlie previously reported variability and provided a framework to enhance reproducibility across studies.

Our SLR identified >60% of variability in MN loss across the three most commonly used SMA mouse models. Given that MN death is a defining hallmark of MN diseases, such variability in this key readout may substantially bias mechanistic interpretation and the evaluation of therapeutic efficacy. The meta-analysis suggested that the examined spinal region is a key source of variability in reported MN loss and that segment–specific upper lumbar and medial MN pools innervating axial/proximal muscles are consistently more affected than lateral MNs innervating distal muscles in SMNΔ7 mice, reflecting the proximo-distal gradient of weakness in SMA patients ^25,26^. To directly test this anatomical selectivity, we validated a sectional ChAT-based quantification method ^20,26,42^ within intact, cleared spinal cords using high-resolution confocal microscopy. This sectional approach yielded precise, segment-specific MN numbers across neonatal and adult mice, establishing a reliable methodology for standardized, cross-study comparisons. The implementation of an automated deep-learning workflow using Cellpose ^51,52^ for reliable MN quantification of intact spinal cords offered an unbiased alternative and further validated this sectional approach.

Using MN counts acquired from applying the sectional approach to the entire lumbar spinal cord in SMNΔ7 mice, we were able to confirm the enhanced vulnerability of proximal muscle-innervating MN pools established in our meta-analysis. Moreover, computational simulations of randomly selecting spinal segments in control and SMNΔ7 mice identified inconsistent rostro–caudal MN populations as a key driver of interstudy variability and showed that proposed stereology-like approaches are unreliable for spinal MN count analyses ^39,62,63^. In contrast, restricting these analyses to MN pools that innervate proximal muscles produced robust, reproducible patterns of degeneration. Together, our established anatomically standardized manual and automated quantification methods provided a platform for MN quantification in health and disease and identified inconsistent spinal segment selection as a major driver of variability in MN numbers.

Although nonspecific spinal segment sampling may help explain most of the observed interstudy variability, additional methodological factors such as spinal cord processing and selection of appropriate marker(s) can affect MN quantification beyond anatomical sampling. Spinal cord extraction by corpectomy preserves segmental integrity and enables reliable segment identification using vertebral or ventral root landmarks ^50,64^. In contrast, hydraulic extrusion damages spinal roots and requires immersion fixation, which compromises immunoreactivity ^62,65^, particularly in adult mice, as we have shown here. While the sectioning method had minimal impact on the reliability of MN quantification and observed loss in SMNΔ7 mice, thicker vibratome sections reduced double-counting and artifacts, especially since the rostro-caudal length of spinal segments changes during development or pathology ^20^.

Beyond sampling and tissue processing, the choice of MN marker influences the accuracy of MN quantification. Retrograde tracer injections provide muscle-specific MN identification ^66^ but are invasive and unreliable when axonal degeneration precedes MN loss, while ventral root backfills are technically demanding and developmentally constrained ^26,67^.

In our study, we show that nearly all back-filled MNs expressed ChAT, confirming it as a highly reliable MN marker in the ventral horn; however, inclusion of ChAT⁺ spinal neurons in the intermediate zone ^38^, together with potential downregulation of ChAT expression under neurodegenerative conditions ^68^, can lead to inaccurate MN quantification. Similar to ChAT, HB9 labeled ∼90% of α- and γ-MNs with minimal off-target labeling as previously described ^54,69^. Whereas SMI-32 is largely confined to proximal neurites ^32,33^ and under-detects MNs, Nissl staining marks nearly all MNs. However, both SMI-32 and Nissl staining also label a large number of non-MNs in the ventral horn, namely spinal interneurons. Our data indicate that these include ventral spinocerebellar tract neurons, Renshaw cells, and FoxP2+ neurons. It is likely that additional ventral interneuron populations, such as V0, V2a/V2b, and V3 cells, may also be misassigned as MNs using SMI-32 and Nissl staining ^70^. Avoiding the possible inclusion of interneurons into MN counts should be taken into great consideration as interneuron vulnerability could vary from MNs and confound results in MN disease mouse models. Within certain disease models, selective shrinkage of MN soma size further exacerbates this limitation ^61,71^, rendering size- or location-based classification with nonspecific markers unreliable. Notably, mislabeling of non-MNs is a less prominent issue in human than murine spinal cords, likely due to the larger size and reduced density of MNs ^72^. Collectively, these results demonstrate that rigorous spinal cord processing and the use of selective MN markers are essential for specific and reliable MN quantification in mice.

Having established a highly reliable MN quantification method in wild-type mice, we applied this approach to reassess regional vulnerability across the spinal cord in the most widely studied mouse models of ALS and SMA ^18,28^. At end-stage in SOD1-G93A mice, α-MN degeneration was uniform across all spinal motor pools, whereas SMNΔ7 mice exhibited selective vulnerability restricted to MNs innervating axial and proximal muscles. These observations suggest that, in SMA more so than in ALS, a substantial subset of MNs is intrinsically resistant to neurodegeneration. Consistent with this notion, autopsy studies of SMA Type I patients have repeatedly reported that ∼50% of MNs are preserved ^73–76^. Although α-MNs are well established as the primary vulnerable population, γ-MNs have generally been considered resistant in both SOD1-G93A and SMA models ^60,61,71^. In concordance, we detected no γ-MN loss in SOD1-G93A mice. Interestingly, spinal segment-specific analysis in SMNΔ7 mice revealed a modest reduction in a subset of γ-MN pools innervating proximal and axial muscles, suggesting that they are not entirely spared but substantially less vulnerable than α-MNs. The extent of this pathology may be underestimated due to the short lifespan of this severe SMA model. Consistent with this interpretation, analyses of SMA patient tissue report γ-MN reductions comparable to α-MN loss ^75^. In contrast to the selective MN loss observed in SMNΔ7 mice, we detected no MN degeneration across the spinal cord in 9-month-old Type III–like SMA mice on a C57BL/6 background. This differs from a previous report describing >30% late-onset loss of MNs innervating distal muscles ^58^, although these MNs are consistently spared across SMA mouse models ^20,26^. Consistent with the absence of MN loss, our study demonstrates that NMJs of both axial and distal muscles remained fully innervated in Type III–like SMA mice, consistent with previously reported preservation of muscle function at this age ^58^. The use of non-specific MN markers may explain conflicting reports from studies of this model on an FVB background, which range from early, non-progressive MN loss to a gradual ∼30% decline by one year of age ^77,78^. Importantly, the severe Taiwanese SMA model, generated by breeding Type III–like SMA mice yet carrying fewer SMN copies, shows no MN loss across both C57BL/6 and FVB backgrounds when assessed using the same validated sectional quantification ^20,26^, indicating that these discrepancies do not stem from genetic backgrounds. Two independent reports using the same approach also demonstrated the absence of MN loss in the milder end-stage Smn^2B/–^ model, although minor, selective MN degeneration emerges beyond median survival ^20,79^. Consistent with this, convergent upregulation and phosphorylation of p53 in MNs drives their degeneration in SMNΔ7 and Smn^2B/−^ mice ^20,42,48,59^, whereas this response is absent in Taiwanese ^20^ and Type III-like SMA mice, linking cell-autonomous p53 activation to MN degeneration across SMA models. Accordingly, the >60% variability in reported MN loss across SMA mouse models identified in our SLR and meta-analysis is likely driven primarily by methodological differences, with robust MN degeneration most consistently observed in the SMNΔ7 model.

In conclusion, our study establishes a standardized, validated, and scalable framework for MN identification and quantification in mouse models across normal development and in the context of injury and neurodegeneration. By reducing methodological variability and enabling reproducible analyses, this approach advances the rigor and translational significance of preclinical studies, ultimately facilitating the development and evaluation of therapeutic strategies for MN disorders.

## Materials and Methods

### Systematic Literature Review

SMA spinal cord studies reporting MN loss between untreated mutant and control littermates were included in the SLR. All relevant studies up to February 23, 2023 were identified in peer-reviewed journals indexed in PubMed using a search strategy described in **Table S1.**

Identified studies were included if they met the pre-specified PICOS criteria adapted for animal studies (population, intervention, comparator, outcomes, and study types; see Table S2). To be included in the SLR, studies (n = 75) had to utilize at least one of three predefined SMA mouse models (SMNΔ7, SMN^2B/–^, or Taiwanese), identified based on breeding origin or genetic model of the mouse; the experimental pairs (mutant and control) had to include at least n = 3; and animals had to be sacrificed within a predetermined, model specific end-stage timeframe, capped by a model-specific maximum postnatal day. MN counts had to be obtained ex vivo from the ventral region of the mouse spinal cord. Any mice receiving treatment were excluded from MN counts. MN count means for study cohorts were extracted from publication text or reliably estimated using the open-source software WebPlotDigitizer ^80^. Exact details for each of the screening criteria are included in Table S3.

### Meta-Analysis

A subset of publications meeting the SLR criteria for the SMNΔ7 model was included in the meta-analysis (43 of 75; **Table S3**). Pooled effect sizes were acquired using random-effects models based on between-group standardized mean differences and corrected for small sample sizes (n < 21) with Hedges’ g. MN count means were pooled and their confidence intervals were obtained either via WebPlotDigitizer or from the figure summaries within the original publications. For studies reporting standard deviations (SD), 95% confidence intervals were calculated from the SDs and corresponding sample sizes. Heterogeneity across studies was quantified via the I² statistic, which estimates the proportion of total variability in effect estimates attributable to between-study heterogeneity rather than sampling error. Higher I² values indicate greater between-study variability, suggesting that differences in study-level characteristics may contribute to the observed outcomes. By contrast, lower I² values indicate more homogeneity across studies, implying that the observed variability is more likely due to chance. A feature of I² is its relative independence from the number of studies included within a meta-analysis. I² is commonly expressed as a percentage and interpreted using conventional thresholds: 0% (none), 25% (low), 50% (medium) and 75% (high) ^81^.

### Subgroup Analysis and Multilevel Analysis

The reported heterogeneity of the meta-analysis was further explored using subgroup or moderator analyses to examine potential sources of between-study heterogeneity. This is done under the premise that not all studies originate from the same population and may differ in their underlying true effects due to study-level characteristics. Groupings of experiments according to hypothesized effect modifiers were tested using a fixed-effects model to assess for differences between subgroups. Within each subgroup, a random-effects model was utilized and a test for heterogeneity was performed. The effects for each subgroup were pooled and compared across subgroups. A change in the I² value was examined descriptively to assess whether the hypothesized variable may contribute to the variation observed between studies. To prevent skewing data towards authors with multiple experiments included in the meta-analysis, multilevel analyses were conducted to account for the non-independence of effect sizes. The R statistical software (Version 4.4.3, R Foundation for Statistical Computing, Vienna, Austria) was utilized for the meta-analysis.

### Mouse lines and genotyping

Breeding and experimental procedures were conducted in the animal facilities of Leipzig University (Leipzig, Saxony, Germany) and Columbia University (New York, NY, USA). All animal work at Leipzig University complied with European regulations (Council Directive 86/609/EEC), the German Animal Welfare Act (Tierschutzgesetz), and was approved by the regional authority (Landesdirektion Leipzig). Experiments at Columbia University were performed in accordance with the U.S. National Institutes of Health Guidelines for the Care and Use of Laboratory Animals and approved by the Columbia University Institutional Animal Care and Use Committee (IACUC). This included a 12h/12h light/dark cycle with unrestricted access to both food and water. The SMNΔ7 mouse model (Smn+/-; SMN2+/+; SMNΔ7+/+) on FVB background was obtained from Jackson Laboratory (stock #005025). The 4-copy SMN2 Type III-like SMA model was originally purchased on a FVB background from the Jackson Laboratory (stock # 005058, FVB.Cg-Tg(SMN2)2HungSmn1tm1Hung/J). Mice were then backcrossed over seven generations to a C57BL/6N background ^82^. For ALS mice, the SOD1-WT B6SJL (JAX stock #002297) and SOD1-G93A B6SJL (JAX stock #002726) lines were used. ChAT-eGFP mice were obtained from the Jackson Laboratory (stock #007902). The following primers were used for genotyping: SMNΔ7: forward sequence (5′ to 3′) = GATGATTCTGACATTTGGGATG, reverse sequences (5′ to 3′) = TGGCTTATCTGGAGTTTCACAA and GAGTAACAACCCGTCGGATTC (wild-type band: 325 bp, mSmn ko: 411 bp). 4-copy SMN2 Type III-like SMA: forward sequences (5′ to 3′) = ATAACACCACCACTCTTACTC and GTAGCCGTGATGCCATTGTCA, reverse sequence (5′ to 3′) = AGCCTGAAGAACGAGATCAGC (wild-type band: 1050 bp, mSmn ko: 950 bp). SOD1-WT and SOD1-G93A: forward sequence (5’ to 3’) = CATCAGCCCTAATCCATCTGA, reverse sequence (5’ to 3’) = CGCGACTAACAATCAAAGTGA (transgene = 236 bp). ChAT-eGFP: forward sequence (5’ to 3’) = AAGTTCATCTGCACCACCG, reverse sequence (5’ to 3’) = CGTCGCCGATGGGGGTGTTC (transgene = 400 bp). For the SMNΔ7 mouse line, control mice were obtained from the same litters as mutant mice. For the 4-copy SMN2 Type III-like SMA line, age-matched C57BL/6N mice were used as controls. For SOD1-G93A mice, age matching controls from the SOD1-WT line were taken. Experimental mice were monitored for welfare in accordance with approved animal protocol guidelines. Similar numbers of male and female mice were included, and data were combined, as no sex-specific differences in MN counts were observed or have been previously reported in SMNΔ7, SOD1-G93A or 4-copy SMN2 Type III-like SMA mice.

### Immunofluorescence of sectioned murine and human spinal cord and murine muscles

Mice were transcardially perfused with phosphate-buffered saline (PBS) followed by 4% paraformaldehyde (PFA), and spinal cords were postfixed overnight in 4% PFA at 4°C. For preparation of non-perfused immersion-fixed spinal cord segments from P10 and P60 ChAT-eGFP mice or P11 SMNΔ7 mice, spinal cords were dissected after animal-welfare-compliant decapitation into ice-cold, oxygenated (95%O_2_–5%CO_2_), and continuously perfused with artificial cerebrospinal fluid (aCSF) containing 113 mM NaCl, 3 mM KCl, 1 mM NaH_2_PO_4_.H_2_O, 25 mM NaHCO_3_, 11 mM D-glucose, 2 mM CaCl_2_.H_2_O and 2 mM MgSO_4_.7H_2_O. After dissection, spinal cords were immediately transferred to 4% PFA for fixation. Fixed tissue was stored in PBS containing 0.01% sodium azide (NaN_3_) until further processing.

Thoracic spinal cord from patients with SMA and controls were collected at Johns Hopkins University School of Medicine, Baltimore, MD, USA, as follows: controls: CNTL #1 (cause of death unknown), CNTL #2 (transverse myelitis; collected and analyzed tissue was not affected by transverse myelitis.), CNTL #3 (cardiac arrest) and CNTL #4 (pneumonia); and as SMA patients: SMA #1, SMA #2, SMA #3, SMA #4, SMA #5. Expedited autopsies were conducted under parental- or patient-informed consent in strict observance of the legal and institutional ethical regulations. Tissues from patients with SMA and age-matched controls with a short post-mortem interval were included. All patient and control tissues used in this study are listed in **Table S4**.

Following spinal cord dissection, mouse spinal cord segments were localized by their ventral roots as described previously ^43^. For vibratome sectioning, individual segments were embedded in 5% agarose and serially sectioned transversely at 75 µm using a Leica VT1000S vibratome. For cryostat sectioning of human and mouse spinal cord tissue, segments were cryoprotected sequentially in 15% and 30% sucrose solutions at 4°C for at least 2 hours and overnight, respectively. The tissue was then embedded in Sakura Tissue-Tek O.C.T. compound and frozen in 2-methylbutane cooled with liquid nitrogen. Serial transverse sections (20 µm) were cut at −20°C on a Leica CM3050 S cryostat and stored at −80°C until use.

Human spinal cord sections were incubated for 20 min in Polyscience L.A.B. solution for antigen retrieval at room temperature and subsequently washed three times in PBS. Human and mouse spinal cord cryostat sections were blocked for 60 minutes at room temperature in PBS containing 5% normal donkey serum and 0.3% Triton X-100 (PBS-T, pH 7.4) and subsequently incubated with primary antibodies for a minimum of 18 hours at room temperature (see **Table S5** for details). SMI-32 antibody was directly conjugated to Alexa 647 and did not require a secondary antibody for visualization. After incubation, tissues were washed three times for 10 minutes each in PBS, then incubated with appropriate fluorophore-conjugated secondary antibodies (Alexa 488, Cy3, Alexa 647, DyLight 405; Jackson ImmunoResearch) diluted 1:1000 in PBS-T for 3 hours at room temperature. Following secondary antibody staining, sections were washed three times for 10 minutes in PBS, mounted on slides, covered with glass coverslips, and protected with an anti-fading solution of glycerol:PBS (3:7), as described previously ^75^.

Mouse spinal cord sections cut with a vibratome were blocked for 90 minutes at room temperature in 5% normal donkey serum (NDS) diluted in PBS-T and incubated with primary antibodies (see **Table S5** for details). Afterwards, the sections were washed six times for 10 minutes each in PBS, then incubated with secondary antibodies diluted 1:1000 in PBS-T for 3 hours at room temperature. Finally, sections were washed six times for 10 minutes in PBS, mounted on slides, covered with glass coverslips, and protected with a solution containing glycerol:PBS (3:7), as described previously ^20^. Labeling of ventral spinocerebellar tract neurons was achieved via injection to the cerebellum of 1% cholera toxin B subunit (CTB) conjugated to Alexa 488 (Invitrogen) in P0 C57BL/6 mice, following established protocols ^31^.

For immunostaining of neuromuscular junctions (NMJs), desired muscles were dissected immediately after perfusion and post-fixed with 4% PFA overnight. After fixation, single muscle fibers were teased apart and washed three times in PBS for 10 min each, followed by incubation for 1 h at room temperature with a blocking solution consisting of PBS-T (0.3%) with 5% NDS. Afterwards mouse anti-neurofilament (NF) and anti-synaptic vesicle 2 (SV2) antibodies to immunolabel the presynaptic aspect of the NMJ were applied in blocking solution overnight at room temperature (**Table S5**). The muscle fibers were then washed three times for 10 min in PBS, followed by incubation with secondary antibodies and α-bungarotoxin (BTX) Alexa Fluor 555 diluted in PBS-T for 3h at room temperature as described previously ^83^. Lastly, muscle fibers were washed three times in PBS for 10 min and mounted on slides covered with glycerol: PBS (3:7).

### Retrograde filling, immunofluorescence and clearing of whole-mount spinal cord

Segment-specific retrograde filling of MNs via ventral roots was performed as established previously ^26^. Briefly, the spinal cord of C57BL/6 P4 mice was dissected free under in vitro conditions in cooled (10°C) and continuously perfused artificial CSF (aCSF). The appropriate ventral roots were inserted into suction electrodes and backfilled in an alternating manner with either Texas Red Dextran or Fluorescein Dextran [10,000 molecular weight (MW)] for 24 h. Following this period, the lumbar spinal cord was immersion-fixed in 4% PFA for 24 h. Spinal cords that were processed for analysis with Cellpose software, were pinned into a custom-made silicone chamber using a 3D-printed mold designed and produced by Juergen Simon, Steffen, and Ingo Rueger. Afterwards, lumbar spinal cords were processed for whole-mount immunostaining and subsequent clearing. All steps were performed in light-protected conditions, on an orbital shaker, in 2 ml Eppendorf microtubes and with solutions containing 0.02% NaN3. Spinal cords were permeabilized for 48 h in PBS with 2% Triton X-100 (2% PBS-T) at room temperature and then blocked for 48 h at 4°C in 1% PBS-T supplemented with 2.5% DMSO and 10% NDS. Next, they were incubated for 4 days at 4°C with ChAT antibody (Millipore) diluted 1:100 in 0.2% PBS-T containing 2.5% DMSO and 1% NDS. After three 1 h washes and an overnight wash in 0.2% PBS-T containing 3% NaCl, samples were incubated for 48 h at 4°C with Alexa 647 anti-goat secondary antibody (Jackson lab) diluted at 1:250 in 0.2% PBS-T containing 2.5% DMSO and 1% NDS. Following identical washing conditions, spinal cords were transferred into RapiClear® 1.49 (SunJinLab) for optical clearing for 24 h at 37°C and glued ventral side down to a glass-bottom dish filled with RapiClear® 1.49. After imaging, spinal cords were removed from the dish and incubated overnight in PBS to reverse clearing. The next day, they were cut at a vibratome into transverse serial sections and mounted directly onto slides in a consecutive manner with PBS:Glycerol (3:7) for final imaging. Similar procedures regarding immunofluorescence and clearing were applied to perfused L1 segments with additional HB9 staining (diluted 1:3000) for whole-mount MN segmentation and counting with Cellpose.

### Confocal microscopy and image analysis

Imaging of tissues was conducted using a Leica SP8, Zeiss LSM 700, or Olympus FV300 inverted confocal microscope. Sectioned spinal cords were imaged using either a 10x objective for hemisection overviews, a 20x objective for MN pool images, with a z-step size of 4 µm between optical planes, or a 63x objective for close-up images. Whole-mount lumbar spinal cords and isolated L1 segments were scanned using a 25x objective at a z-step size of 1 µm. NMJs were imaged using a 20x objective and a z-step size of 4 µm.

For retrogradely filled spinal cords, segment borders were determined from retrograde labeling for both whole-mount imaging and after sectioning. Only ChAT+ MNs located in the ventral horn and containing a visible nucleus were counted, to avoid double-counting from adjoining sections. For cryostat-sectioned spinal cord segments, every third section was analyzed to further prevent duplicate counting of the same MNs. When applicable, NeuN+/ChAT+ MNs were classified as α-MNs, whereas ChAT+ MNs lacking NeuN immunoreactivity were classified as γ-MNs. If stated, Abercrombie correction was applied the following way: 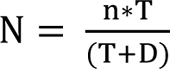 with N = corrected number of MNs, n = original counted number of MNs, T = section thickness in µm, D = mean diameter of MN nucleus in µm.

For analysis of the other MN markers HB9, Nissl and SMI-32 in comparison to ChAT, confocal images were acquired with a 10x objective. MN pools of each analyzed slice were measured in length and width according to ChAT signaling and the mean plus 3 times the SD width and length was calculated from the data acquired from 5 slices within each mouse. This calculated width and length were then applied to the corresponding MN pools for analysis. To guarantee that all MNs were counted, we used the previously reported mean γ-MN area (232.4 µm²; ^53^) and subtracted three SD, yielding 82.4 µm², with all marker-positive signals exceeding this area included in the analysis. The amount of marker+ MNs (ChAT+) was calculated as well as the amount of putative MNs that were ChAT-but marker+.

To quantify MN soma size, the soma perimeter was outlined in the z-plane showing the largest nuclear cross-section, and the enclosed area was measured in µm². At least five sections were analyzed per animal, or 3 for human spinal cords. For all analyses, either Leica Las X software or Fiji Image-J software was used.

### Training and implementation of MN quantification with Cellpose software

Confocal z-stack images of whole-mount L1 spinal cord segments from control and SMNΔ7 mice, immunolabeled for ChAT and HB9, were preprocessed in FIJI (ImageJ) to optimize Cellpose-based counting. For the HB9 channel, 3D stacks were despeckled, median-filtered (radius 0.3 pixels), background-subtracted (rolling ball radius 60 pixels), and histogram-equalized to normalize contrast. For ChAT, brightness and contrast were adjusted. The TissueNet model in Cellpose was chosen as the base and further trained with human-in-the-loop annotation using five representative 2D images from both control and SMNΔ7 samples^51,52^.

After training, the optimized model was applied to the corresponding 3D stacks to segment ChAT+ MNs using HB9 nuclear signal as an auxiliary marker and generate automated counts. These automated MN counts were directly compared to manual counts obtained by manual visual inspection of the same image stacks. Segmentation served solely as an intermediate detection step for enumeration; no pixel-level segmentation accuracy was assessed, as the aim was to compare MN counts between the two methods. To quantify correspondence between automated and manual counts, precision was calculated as: 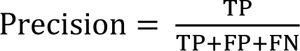 where true positives (TP) were MNs identified by both manual counting and Cellpose segmentation, false positives (FP) were MNs segmented by Cellpose but not counted manually, and false negatives (FN) were MNs counted manually but missed by Cellpose. This definition of precision is adapted here to reflect agreement in MN counting rather than classical pixel-wise segmentation performance metrics.

### Randomization approach for sampling from spinal segments

MN counts per vibratome section for individual lumbar segments (L1–L6) were obtained from n = 4 control and SMNΔ7 mice, as described above. These datasets were used as input for a custom R script (R Foundation for Statistical Computing, Vienna, Austria) to evaluate how different randomized sampling strategies, similar to those commonly applied in the field, affect the detection of MN loss. In each simulation run, sampling points were randomly selected to simulate the following approaches: drawing five consecutive sections from anywhere in the lumbar region, selecting every 2^nd^ or every 10^th^ section through the entire lumbar cord starting at a random position in L1, or selecting five random sections within L1 or L2 separately. The R script was executed four times for each sampling scheme, with each execution generating a new independent random selection (“simulation round”). For every round, the resulting subsets from the control and SMNΔ7 datasets were used to calculate mean MN counts per genotype, and the values were statistically compared to determine whether each strategy could detect significant MN loss.

### Quantification and statistical analysis

All experimental results are expressed using a calculated mean and bounded by the standard error of the mean (SEM). In this manuscript, n refers to the number of patients or mice used in each group. Each experiment included at least three biological replicates per group. To assess the symmetry of data distribution, the Shapiro-Wilk normality test was applied. For comparisons between two groups, statistical analysis depended on the data’s distribution and the equality of standard deviations (SDs). If the data followed a parametric distribution and the SDs were equal, an unpaired t-test was used. When the SDs were unequal, Welch’s t-test was applied. For nonparametric distributions, the Mann-Whitney test was performed. For comparisons among three groups or more, a one-way ANOVA followed by Tukey’s multiple comparison test was used for parametric distributions. Nonparametric distributions were analyzed using the Kruskal-Wallis test with Dunn’s correction. When data was compared against a set value and the distribution was parametric, a one-sample t-test, when the distribution was non-parametric, a Wilcoxon signed-ranked test was performed. The statistical tests used for each experiment are indicated in the respective figure legends. All statistical analyses were conducted using GraphPad Prism 10, and p-values are reported within the figures.

## Data availability

The authors confirm that the data supporting the findings of this study are available within the article and its Supplementary material. Some data are not publicly available owing to patient-related restrictions, because they contain information that could compromise the privacy of research participants. Additionally, certain derived data from mouse experiments are available from the corresponding authors upon reasonable request.

## Acknowledgement

We thank Drs. Livio Pellizzoni and Emily Lowry for their critical comments on the manuscript. We are grateful to Juergen Simon, Ingo Rueger, and Steffen Rueger for designing and producing the 3D-printed mold. We also thank Susan Brenner-Morton and Dr. Hynek Wichterle for providing the HB9 antibody. We acknowledge Dr. Daniel Lin from SUNJIN Lab Co. for providing RapiClear solutions, as well as technical assistance and protocols. We thank Dr. Anja Reinert for introducing us to the Cellpose framework. Finally, we are grateful to Drs. Stefan Hallermann, Johannes Hirrlinger, Jens Eilers, and Tobias Langenhan for providing reagents and access to facilities. During the peer review process, Biogen had the opportunity to review the manuscript. The authors had full editorial control of the manuscript and provided their final approval on all content.

## References

1. Stifani, N. (2014). Motor neurons and the generation of spinal motor neuron diversity. Front Cell Neurosci 8, 293. 10.3389/fncel.2014.00293.

2. Chaudhary, R., Agarwal, V., Rehman, M., Kaushik, A.S., and Mishra, V. (2022). Genetic architecture of motor neuron diseases. J Neurol Sci 434, 120099. 10.1016/j.jns.2021.120099.

3. Orrell, R.W. (2010). Motor neuron disease: systematic reviews of treatment for ALS and SMA. Br Med Bull 93, 145–159. 10.1093/bmb/ldp049.

4. Yaginuma, H., Sato, N., Homma, S., and Oppenheim, R.W. (2001). Roles of caspases in the programmed cell death of motoneurons in vivo. Arch Histol Cytol 64, 461–474. 10.1679/aohc.64.461.

5. Houenou, L.J., Li, L., Lo, A.C., Yan, Q., and Oppenheim, R.W. (1994). Naturally occurring and axotomy-induced motoneuron death and its prevention by neurotrophic agents: a comparison between chick and mouse. Prog Brain Res 102, 217–226. 10.1016/S0079-6123(08)60542-7.

6. Celik, M., Gokmen, N., Erbayraktar, S., Akhisaroglu, M., Konakc, S., Ulukus, C., Genc, S., Genc, K., Sagiroglu, E., Cerami, A., and Brines, M. (2002). Erythropoietin prevents motor neuron apoptosis and neurologic disability in experimental spinal cord ischemic injury. Proc Natl Acad Sci U S A 99, 2258–2263. 10.1073/pnas.042693799.

7. Prow, N.A., and Irani, D.N. (2008). The inflammatory cytokine, interleukin-1 beta, mediates loss of astroglial glutamate transport and drives excitotoxic motor neuron injury in the spinal cord during acute viral encephalomyelitis. J Neurochem 105, 1276–1286. 10.1111/j.1471-4159.2008.05230.x.

8. Li, L., Oppenheim, R.W., Lei, M., and Houenou, L.J. (1994). Neurotrophic agents prevent motoneuron death following sciatic nerve section in the neonatal mouse. J Neurobiol 25, 759–766. 10.1002/neu.480250702.

9. Xu, W., Chi, L., Xu, R., Ke, Y., Luo, C., Cai, J., Qiu, M., Gozal, D., and Liu, R. (2005). Increased production of reactive oxygen species contributes to motor neuron death in a compression mouse model of spinal cord injury. Spinal Cord 43, 204–213. 10.1038/sj.sc.3101674.

10. Liewluck, T., and Saperstein, D.S. (2015). Progressive Muscular Atrophy. Neurol Clin 33, 761–773. 10.1016/j.ncl.2015.07.005.

11. Hardiman, O., Al-Chalabi, A., Chio, A., Corr, E.M., Logroscino, G., Robberecht, W., Shaw, P.J., Simmons, Z., and van den Berg, L.H. (2017). Amyotrophic lateral sclerosis. Nat Rev Dis Primers 3, 17071. 10.1038/nrdp.2017.71.

12. Breza, M., and Koutsis, G. (2019). Kennedy’s disease (spinal and bulbar muscular atrophy): a clinically oriented review of a rare disease. J Neurol 266, 565–573. 10.1007/s00415-018-8968-7.

13. Jablonka, S., and Yildirim, E. (2024). Disease Mechanisms and Therapeutic Approaches in SMARD1-Insights from Animal Models and Cell Models. Biomedicines 12. 10.3390/biomedicines12040845.

14. Wirth, B. (2021). Spinal Muscular Atrophy: In the Challenge Lies a Solution. Trends Neurosci 44, 306–322. 10.1016/j.tins.2020.11.009.

15. Tsai, L.K., Tsai, M.S., Ting, C.H., and Li, H. (2008). Multiple therapeutic effects of valproic acid in spinal muscular atrophy model mice. J Mol Med (Berl) 86, 1243–1254. 10.1007/s00109-008-0388-1.

16. Mattis, V.B., Ebert, A.D., Fosso, M.Y., Chang, C.W., and Lorson, C.L. (2009). Delivery of a read-through inducing compound, TC007, lessens the severity of a spinal muscular atrophy animal model. Hum Mol Genet 18, 3906–3913. 10.1093/hmg/ddp333.

17. Cervero, C., Blasco, A., Tarabal, O., Casanovas, A., Piedrafita, L., Navarro, X., Esquerda, J.E., and Caldero, J. (2018). Glial Activation and Central Synapse Loss, but Not Motoneuron Degeneration, Are Prevented by the Sigma-1 Receptor Agonist PRE-084 in the Smn2B/-Mouse Model of Spinal Muscular Atrophy. J Neuropathol Exp Neurol 77, 577–597. 10.1093/jnen/nly033.

18. Le, T.T., Pham, L.T., Butchbach, M.E., Zhang, H.L., Monani, U.R., Coovert, D.D., Gavrilina, T.O., Xing, L., Bassell, G.J., and Burghes, A.H. (2005). SMNDelta7, the major product of the centromeric survival motor neuron (SMN2) gene, extends survival in mice with spinal muscular atrophy and associates with full-length SMN. Hum Mol Genet 14, 845–857. 10.1093/hmg/ddi078.

19. Eshraghi, M., McFall, E., Gibeault, S., and Kothary, R. (2016). Effect of genetic background on the phenotype of the Smn2B/-mouse model of spinal muscular atrophy. Hum Mol Genet 25, 4494–4506. 10.1093/hmg/ddw278.

20. Buettner, J.M., Sime Longang, J.K., Gerstner, F., Apel, K.S., Blanco-Redondo, B., Sowoidnich, L., Janzen, E., Langenhan, T., Wirth, B., and Simon, C.M. (2021). Central synaptopathy is the most conserved feature of motor circuit pathology across spinal muscular atrophy mouse models. iScience 24, 103376. 10.1016/j.isci.2021.103376.

21. Charles Watson, G.P., Gulgun Kayalioglu, Claire Heise, (2009). The Spinal Cord. Academic Press Chapter 16 - Atlas of the Mouse Spinal Cord, Pages 308–379. 10.1016/B978-0-12-374247-6.50020-1.

22. Dasen, J.S., De Camilli, A., Wang, B., Tucker, P.W., and Jessell, T.M. (2008). Hox repertoires for motor neuron diversity and connectivity gated by a single accessory factor, FoxP1. Cell 134, 304–316. 10.1016/j.cell.2008.06.019.

23. Dasen, J.S., Tice, B.C., Brenner-Morton, S., and Jessell, T.M. (2005). A Hox regulatory network establishes motor neuron pool identity and target-muscle connectivity. Cell 123, 477–491. 10.1016/j.cell.2005.09.009.

24. Demireva, E.Y., Shapiro, L.S., Jessell, T.M., and Zampieri, N. (2011). Motor neuron position and topographic order imposed by beta- and gamma-catenin activities. Cell 147, 641–652. 10.1016/j.cell.2011.09.037.

25. Lee, J.C., Chung, W.K., Pisapia, D.J., and Henderson, C.E. (2025). Motor pool selectivity of neuromuscular degeneration in type I spinal muscular atrophy is conserved between human and mouse. Hum Mol Genet 34, 347–367. 10.1093/hmg/ddae190.

26. Mentis, G.Z., Blivis, D., Liu, W., Drobac, E., Crowder, M.E., Kong, L., Alvarez, F.J., Sumner, C.J., and O’Donovan, M.J. (2011). Early functional impairment of sensory-motor connectivity in a mouse model of spinal muscular atrophy. Neuron 69, 453–467. 10.1016/j.neuron.2010.12.032.

27. Simon, N.G., Lee, M., Bae, J.S., Mioshi, E., Lin, C.S., Pfluger, C.M., Henderson, R.D., Vucic, S., Swash, M., Burke, D., and Kiernan, M.C. (2015). Dissociated lower limb muscle involvement in amyotrophic lateral sclerosis. J Neurol 262, 1424–1432. 10.1007/s00415-015-7721-8.

28. Gurney, M.E., Pu, H., Chiu, A.Y., Dal Canto, M.C., Polchow, C.Y., Alexander, D.D., Caliendo, J., Hentati, A., Kwon, Y.W., Deng, H.X., and, et al. (1994). Motor neuron degeneration in mice that express a human Cu,Zn superoxide dismutase mutation. Science 264, 1772–1775. 10.1126/science.8209258.

29. Bowerman, M., Beauvais, A., Anderson, C.L., and Kothary, R. (2010). Rho-kinase inactivation prolongs survival of an intermediate SMA mouse model. Hum Mol Genet 19, 1468–1478. 10.1093/hmg/ddq021.

30. Guo, J., Qiu, W., Soh, S.L., Wei, S., Radda, G.K., Ong, W.Y., Pang, Z.P., and Han, W. (2013). Motor neuron degeneration in a mouse model of seipinopathy. Cell Death Dis 4, e535. 10.1038/cddis.2013.64.

31. Chalif, J.I., Martinez-Silva, M.L., Pagiazitis, J.G., Murray, A.J., and Mentis, G.Z. (2022). Control of mammalian locomotion by ventral spinocerebellar tract neurons. Cell 185, 328–344 e326. 10.1016/j.cell.2021.12.014.

32. Carriedo, S.G., Yin, H.Z., and Weiss, J.H. (1996). Motor neurons are selectively vulnerable to AMPA/kainate receptor-mediated injury in vitro. J Neurosci 16, 4069–4079. 10.1523/JNEUROSCI.16-13-04069.1996.

33. Tsang, Y.M., Chiong, F., Kuznetsov, D., Kasarskis, E., and Geula, C. (2000). Motor neurons are rich in non-phosphorylated neurofilaments: cross-species comparison and alterations in ALS. Brain Res 861, 45–58. 10.1016/s0006-8993(00)01954-5.

34. Biagioni, F., Ferrucci, M., Ryskalin, L., Fulceri, F., Lazzeri, G., Calierno, M.T., Busceti, C.L., Ruffoli, R., and Fornai, F. (2017). Protective effects of long-term lithium administration in a slowly progressive SMA mouse model. Arch Ital Biol 155, 118–130. 10.12871/00039829201749.

35. Borges, L.F., and Iversen, S.D. (1986). Topography of choline acetyltransferase immunoreactive neurons and fibers in the rat spinal cord. Brain Res 362, 140–148. 10.1016/0006-8993(86)91407-1.

36. Wilson, J.M., Hartley, R., Maxwell, D.J., Todd, A.J., Lieberam, I., Kaltschmidt, J.A., Yoshida, Y., Jessell, T.M., and Brownstone, R.M. (2005). Conditional rhythmicity of ventral spinal interneurons defined by expression of the Hb9 homeodomain protein. J Neurosci 25, 5710–5719. 10.1523/JNEUROSCI.0274-05.2005.

37. Arber, S., Han, B., Mendelsohn, M., Smith, M., Jessell, T.M., and Sockanathan, S. (1999). Requirement for the homeobox gene Hb9 in the consolidation of motor neuron identity. Neuron 23, 659–674. 10.1016/s0896-6273(01)80026-x.

38. Mecca, J., Mignot, J., Gervais, M., Ozturk, T., Astord, S., Berthier, J., Bauche, S., Messeant, J., Biferi, M.G., Rouard, H., et al. (2026). Targeted knockdown of Smn in muscle stem cells induces non-cell autonomous loss of motor neurons. Brain. 10.1093/brain/awag045.

39. Ferrucci, M., Lazzeri, G., Flaibani, M., Biagioni, F., Cantini, F., Madonna, M., Bucci, D., Limanaqi, F., Soldani, P., and Fornai, F. (2018). In search for a gold-standard procedure to count motor neurons in the spinal cord. Histol Histopathol 33, 1021–1046. 10.14670/HH-11-983.

40. Bacskai, T., Rusznak, Z., Paxinos, G., and Watson, C. (2014). Musculotopic organization of the motor neurons supplying the mouse hindlimb muscles: a quantitative study using Fluoro-Gold retrograde tracing. Brain Struct Funct 219, 303–321. 10.1007/s00429-012-0501-7.

41. Fletcher, E.V., Simon, C.M., Pagiazitis, J.G., Chalif, J.I., Vukojicic, A., Drobac, E., Wang, X., and Mentis, G.Z. (2017). Reduced sensory synaptic excitation impairs motor neuron function via Kv2.1 in spinal muscular atrophy. Nat Neurosci 20, 905–916. 10.1038/nn.4561.

42. Simon, C.M., Dai, Y., Van Alstyne, M., Koutsioumpa, C., Pagiazitis, J.G., Chalif, J.I., Wang, X., Rabinowitz, J.E., Henderson, C.E., Pellizzoni, L., and Mentis, G.Z. (2017). Converging Mechanisms of p53 Activation Drive Motor Neuron Degeneration in Spinal Muscular Atrophy. Cell Rep 21, 3767–3780. 10.1016/j.celrep.2017.12.003.

43. Buettner, J.M., Kirmann, T., Mentis, G.Z., Hallermann, S., and Simon, C.M. (2022). Laser microscopy acquisition and analysis of premotor synapses in the murine spinal cord. STAR Protoc 3, 101236. 10.1016/j.xpro.2022.101236.

44. Bikoff, J.B., Gabitto, M.I., Rivard, A.F., Drobac, E., Machado, T.A., Miri, A., Brenner-Morton, S., Famojure, E., Diaz, C., Alvarez, F.J., et al. (2016). Spinal Inhibitory Interneuron Diversity Delineates Variant Motor Microcircuits. Cell 165, 207–219. 10.1016/j.cell.2016.01.027.

45. Floyd, T.L., Dai, Y., and Ladle, D.R. (2018). Characterization of calbindin D28k expressing interneurons in the ventral horn of the mouse spinal cord. Dev Dyn 247, 185–193. 10.1002/dvdy.24601.

46. Abercrombie, M. (1946). Estimation of nuclear population from microtome sections. Anat Rec 94, 239–247. 10.1002/ar.1090940210.

47. Simon, C.M., Blanco-Redondo, B., Buettner, J.M., Pagiazitis, J.G., Fletcher, E.V., Sime Longang, J.K., and Mentis, G.Z. (2021). Chronic Pharmacological Increase of Neuronal Activity Improves Sensory-Motor Dysfunction in Spinal Muscular Atrophy Mice. J Neurosci 41, 376–389. 10.1523/JNEUROSCI.2142-20.2020.

48. Simon, C.M., Van Alstyne, M., Lotti, F., Bianchetti, E., Tisdale, S., Watterson, D.M., Mentis, G.Z., and Pellizzoni, L. (2019). Stasimon Contributes to the Loss of Sensory Synapses and Motor Neuron Death in a Mouse Model of Spinal Muscular Atrophy. Cell Rep 29, 3885–3901 e3885. 10.1016/j.celrep.2019.11.058.

49. Gerstner, F., Wittig, S., Menedo, C., Ruwald, S., Carlini, M.J., Vankova, A., Sowoidnich, L., Martin-Lopez, G., Dreilich, V., Alonso Collado, A., et al. (2025). Cerebellar pathology contributes to neurodevelopmental deficits in spinal muscular atrophy. Brain. 10.1093/brain/awaf336.

50. Buettner, J.M., Sowoidnich, L., Gerstner, F., Blanco-Redondo, B., Hallermann, S., and Simon, C.M. (2022). p53-dependent c-Fos expression is a marker but not executor for motor neuron death in spinal muscular atrophy mouse models. Front Cell Neurosci 16, 1038276. 10.3389/fncel.2022.1038276.

51. Stringer, C., Wang, T., Michaelos, M., and Pachitariu, M. (2021). Cellpose: a generalist algorithm for cellular segmentation. Nat Methods 18, 100–106. 10.1038/s41592-020-01018-x.

52. Pachitariu, M., and Stringer, C. (2022). Cellpose 2.0: how to train your own model. Nat Methods 19, 1634–1641. 10.1038/s41592-022-01663-4.

53. Friese, A., Kaltschmidt, J.A., Ladle, D.R., Sigrist, M., Jessell, T.M., and Arber, S. (2009). Gamma and alpha motor neurons distinguished by expression of transcription factor Err3. Proc Natl Acad Sci U S A 106, 13588–13593. 10.1073/pnas.0906809106.

54. Shneider, N.A., Brown, M.N., Smith, C.A., Pickel, J., and Alvarez, F.J. (2009). Gamma motor neurons express distinct genetic markers at birth and require muscle spindle-derived GDNF for postnatal survival. Neural Dev 4, 42. 10.1186/1749-8104-4-42.

55. Piras, A., Schiaffino, L., Boido, M., Valsecchi, V., Guglielmotto, M., De Amicis, E., Puyal, J., Garcera, A., Tamagno, E., Soler, R.M., and Vercelli, A. (2017). Inhibition of autophagy delays motoneuron degeneration and extends lifespan in a mouse model of spinal muscular atrophy. Cell Death Dis 8, 3223. 10.1038/s41419-017-0086-4.

56. Cervero, C., Montull, N., Tarabal, O., Piedrafita, L., Esquerda, J.E., and Caldero, J. (2016). Chronic Treatment with the AMPK Agonist AICAR Prevents Skeletal Muscle Pathology but Fails to Improve Clinical Outcome in a Mouse Model of Severe Spinal Muscular Atrophy. Neurotherapeutics 13, 198–216. 10.1007/s13311-015-0399-x.

57. Abera, M.B., Xiao, J., Nofziger, J., Titus, S., Southall, N., Zheng, W., Moritz, K.E., Ferrer, M., Cherry, J.J., Androphy, E.J., et al. (2016). ML372 blocks SMN ubiquitination and improves spinal muscular atrophy pathology in mice. JCI Insight 1, e88427. 10.1172/jci.insight.88427.

58. Tsai, L.K., Tsai, M.S., Lin, T.B., Hwu, W.L., and Li, H. (2006). Establishing a standardized therapeutic testing protocol for spinal muscular atrophy. Neurobiol Dis 24, 286–295. 10.1016/j.nbd.2006.07.004.

59. Van Alstyne, M., Simon, C.M., Sardi, S.P., Shihabuddin, L.S., Mentis, G.Z., and Pellizzoni, L. (2018). Dysregulation of Mdm2 and Mdm4 alternative splicing underlies motor neuron death in spinal muscular atrophy. Genes Dev 32, 1045–1059. 10.1101/gad.316059.118.

60. McLeod, V.M., Chiam, M.D.F., Perera, N.D., Lau, C.L., Boon, W.C., and Turner, B.J. (2022). Mapping Motor Neuron Vulnerability in the Neuraxis of Male SOD1(G93A) Mice Reveals Widespread Loss of Androgen Receptor Occurring Early in Spinal Motor Neurons. Front Endocrinol (Lausanne) 13, 808479. 10.3389/fendo.2022.808479.

61. Lalancette-Hebert, M., Sharma, A., Lyashchenko, A.K., and Shneider, N.A. (2016). Gamma motor neurons survive and exacerbate alpha motor neuron degeneration in ALS. Proc Natl Acad Sci U S A 113, E8316–E8325. 10.1073/pnas.1605210113.

62. Austin, A., Beresford, L., Price, G., Cunningham, T., Kalmar, B., and Yon, M. (2022). Sectioning and Counting of Motor Neurons in the L3 to L6 Region of the Adult Mouse Spinal Cord. Curr Protoc 2, e428. 10.1002/cpz1.428.

63. d’Errico, P., Boido, M., Piras, A., Valsecchi, V., De Amicis, E., Locatelli, D., Capra, S., Vagni, F., Vercelli, A., and Battaglia, G. (2013). Selective vulnerability of spinal and cortical motor neuron subpopulations in delta7 SMA mice. PLoS One 8, e82654. 10.1371/journal.pone.0082654.

64. Harrison, M., O’Brien, A., Adams, L., Cowin, G., Ruitenberg, M.J., Sengul, G., and Watson, C. (2013). Vertebral landmarks for the identification of spinal cord segments in the mouse. Neuroimage 68, 22–29. 10.1016/j.neuroimage.2012.11.048.

65. Richner, M., Jager, S.B., Siupka, P., and Vaegter, C.B. (2017). Hydraulic Extrusion of the Spinal Cord and Isolation of Dorsal Root Ganglia in Rodents. J Vis Exp. 10.3791/55226.

66. Mendelsohn, A.I., Simon, C.M., Abbott, L.F., Mentis, G.Z., and Jessell, T.M. (2015). Activity Regulates the Incidence of Heteronymous Sensory-Motor Connections. Neuron 87, 111–123. 10.1016/j.neuron.2015.05.045.

67. Blivis, D., Falgairolle, M., and O’Donovan, M.J. (2019). Dye-coupling between neonatal spinal motoneurons and interneurons revealed by prolonged back-filling of a ventral root with a low molecular weight tracer in the mouse. Sci Rep 9, 3201. 10.1038/s41598-019-39881-0.

68. Ramirez-Jarquin, U.N., Lazo-Gomez, R., Tovar, Y.R.L.B., and Tapia, R. (2014). Spinal inhibitory circuits and their role in motor neuron degeneration. Neuropharmacology 82, 101–107. 10.1016/j.neuropharm.2013.10.003.

69. Koronfel, L.M., Kanning, K.C., Alcos, A., Henderson, C.E., and Brownstone, R.M. (2021). Elimination of glutamatergic transmission from Hb9 interneurons does not impact treadmill locomotion. Sci Rep 11, 16008. 10.1038/s41598-021-95143-y.

70. Brownstone, R.M., and Bui, T.V. (2010). Spinal interneurons providing input to the final common path during locomotion. Prog Brain Res 187, 81–95. 10.1016/B978-0-444-53613-6.00006-X.

71. Powis, R.A., and Gillingwater, T.H. (2016). Selective loss of alpha motor neurons with sparing of gamma motor neurons and spinal cord cholinergic neurons in a mouse model of spinal muscular atrophy. J Anat 228, 443–451. 10.1111/joa.12419.

72. Bahney, J., and von Bartheld, C.S. (2018). The Cellular Composition and Glia-Neuron Ratio in the Spinal Cord of a Human and a Nonhuman Primate: Comparison With Other Species and Brain Regions. Anat Rec (Hoboken) 301, 697–710. 10.1002/ar.23728.

73. Allardyce, H., Lawrence, B.D., Crawford, T.O., Sumner, C.J., and Parson, S.H. (2024). A reassessment of spinal cord pathology in severe infantile spinal muscular atrophy: Reassessment of spinal cord pathology. Neuropathol Appl Neurobiol 50, e13013. 10.1111/nan.13013.

74. Kumagai, T., and Hashizume, Y. (1982). Morphological and morphometric studies on the spinal cord lesion in Werdnig-Hoffmann disease. Brain Dev 4, 87–96. 10.1016/s0387-7604(82)80002-8.

75. Simon, C.M., Delestree, N., Montes, J., Sowoidnich, L., Gerstner, F., Carranza, E., Buettner, J.M., Pagiazitis, J.G., Prat-Ortega, G., Ensel, S., et al. (2025). Proprioceptive synaptic dysfunction is a key feature in mice and humans with spinal muscular atrophy. Brain. 10.1093/brain/awaf074.

76. Kong, L., Valdivia, D.O., Simon, C.M., Hassinan, C.W., Delestree, N., Ramos, D.M., Park, J.H., Pilato, C.M., Xu, X., Crowder, M., et al. (2021). Impaired prenatal motor axon development necessitates early therapeutic intervention in severe SMA. Sci Transl Med 13. 10.1126/scitranslmed.abb6871.

77. Liu, H.C., Ting, C.H., Wen, H.L., Tsai, L.K., Hsieh-Li, H.M., Li, H., and Lin-Chao, S. (2013). Sodium vanadate combined with L-ascorbic acid delays disease progression, enhances motor performance, and ameliorates muscle atrophy and weakness in mice with spinal muscular atrophy. BMC Med 11, 38. 10.1186/1741-7015-11-38.

78. Neuhoff, S., Schmitt, L.I., Liebig, K.C., Hezel, S., Tilahun, N.I., Kleinschnitz, C., Leo, M., and Hagenacker, T. (2025). Neuronal Pentraxin 2 as a Potential Biomarker for Nusinersen Therapy Response in Adults with Spinal Muscular Atrophy: A Pilot Study. Biomedicines 13. 10.3390/biomedicines13081821.

79. Carlini, M.J., Triplett, M.K., and Pellizzoni, L. (2022). Neuromuscular denervation and deafferentation but not motor neuron death are disease features in the Smn2B/-mouse model of SMA. PLoS One 17, e0267990. 10.1371/journal.pone.0267990.

80. Drevon, D., Fursa, S.R., and Malcolm, A.L. (2017). Intercoder Reliability and Validity of WebPlotDigitizer in Extracting Graphed Data. Behav Modif 41, 323–339. 10.1177/0145445516673998.

81. Higgins, J.P., Thompson, S.G., Deeks, J.J., and Altman, D.G. (2003). Measuring inconsistency in meta-analyses. BMJ 327, 557–560. 10.1136/bmj.327.7414.557.

82. Ackermann, B., Krober, S., Torres-Benito, L., Borgmann, A., Peters, M., Hosseini Barkooie, S.M., Tejero, R., Jakubik, M., Schreml, J., Milbradt, J., et al. (2013). Plastin 3 ameliorates spinal muscular atrophy via delayed axon pruning and improves neuromuscular junction functionality. Hum Mol Genet 22, 1328–1347. 10.1093/hmg/dds540.

83. Simon, C.M., Jablonka, S., Ruiz, R., Tabares, L., and Sendtner, M. (2010). Ciliary neurotrophic factor-induced sprouting preserves motor function in a mouse model of mild spinal muscular atrophy. Hum Mol Genet 19, 973–986. 10.1093/hmg/ddp562.

