## Supplemental Data for "A standardized framework resolves ambiguity in motor neuron loss across neurodegenerative diseases"

A

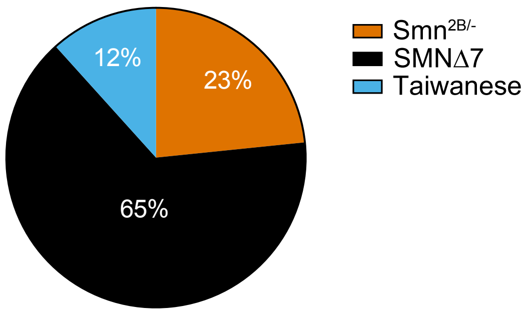


B

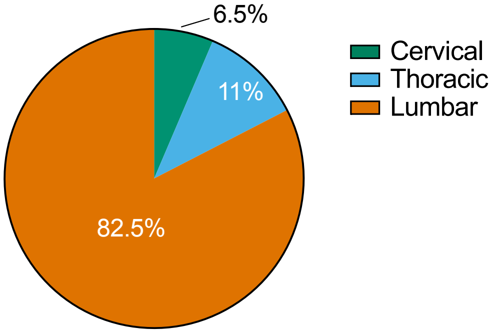


C

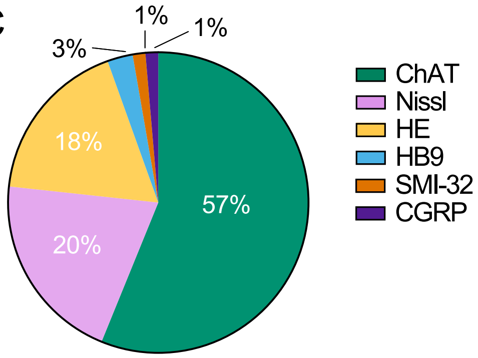


D

| Comparison | ESMD | CI | p-value |
|---|---|---|---|
| L3-L6 vs L5 Med | 0.92 | [-0.35,2.18] | 0.152 |
| L3-L6 vs Lumbar | 0.16 | [-0.80,1.12] | 0.734 |
| L5 Med vs Lumbar | -0.76 | [-1.99,0.46] | 0.221 |
| L3-L6 vs L1-L2 | 2.00 | [1.03,2.98] | <0.0001 |
| L5 Med vs L1-L2 | 1.08 | [-0.14,2.30] | 0.0828 |
| Lumbar vs L1-L2 | 1.84 | [0.97,2.70] | <0.0001 |

E

| Comparison | PMD | CrI | PP | ER |
|---|---|---|---|---|
| L3-L6 vs L5 Med | 0.78 | [-0.35,1.91] | 0.665 | 1.98 |
| L3-L6 vs Lumbar | 0.23 | [-0.54,1.00] | 0.860 | 6.16 |
| L5 Med vs Lumbar | -0.56 | [-1.61,0.49] | 0.764 | 3.24 |
| L3-L6 vs L1-L2 | 1.90 | [0.90,2.91] | 0.992 | 124 |
| L5 Med vs L1-L2 | 1.12 | [-0.12,2.34] | 0.494 | 0.976 |
| Lumbar vs L1-L2 | 1.67 | [0.77,2.58] | 0.989 | 91 |

F

| Comparison | LogLik | AIC | BIC |
|---|---|---|---|
| Multilevel Model | -95.60 | 197.20 | 203.11 |
| Subgroup Model | -82.74 | 177.48 | 188.95 |

Suppl. Figure 1

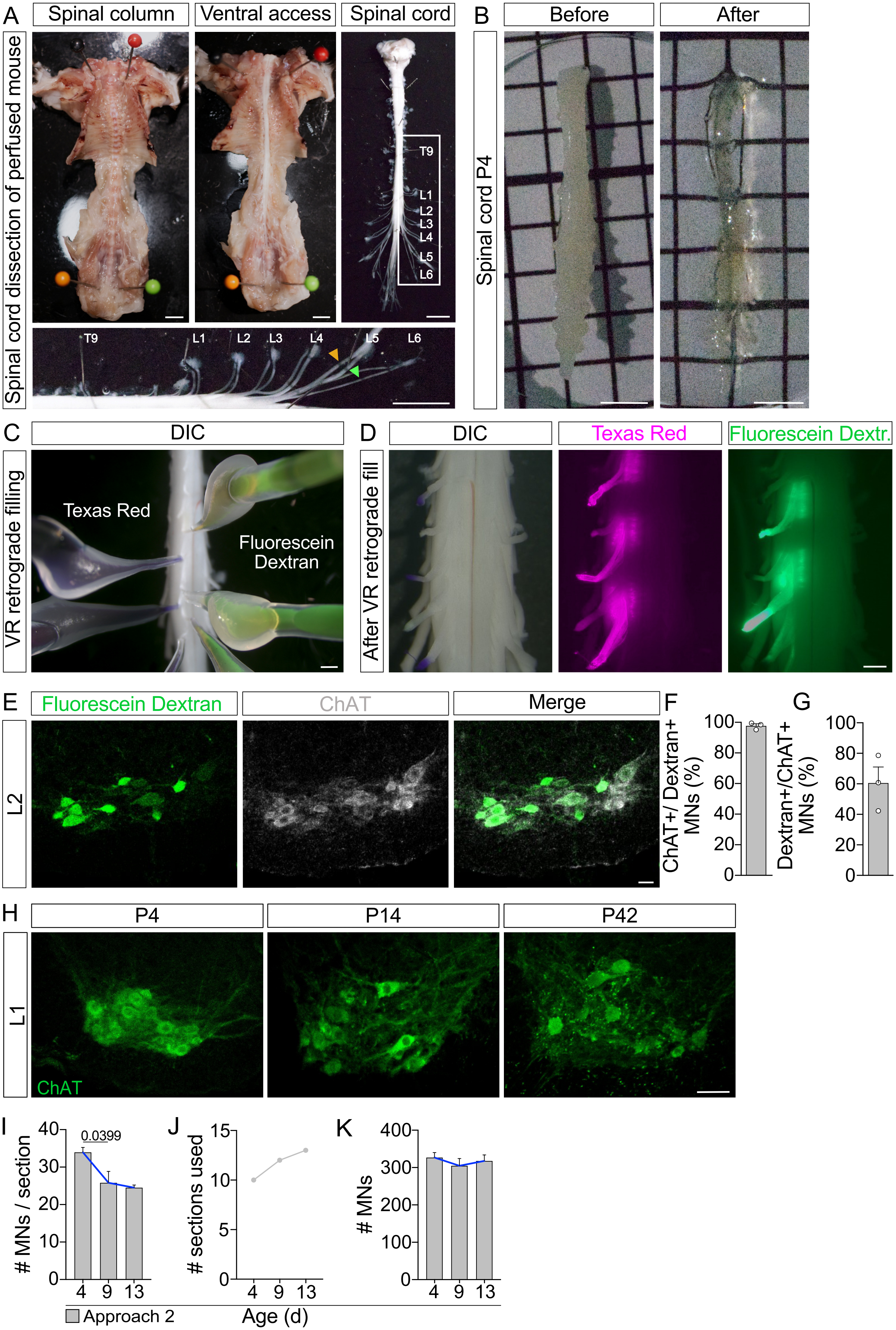

A
Spinal column
Ventral access
Spinal cord
Spinal cord dissection of perfused mouse
T9
L1
L2
L3
L4
L5
L6
B
Before
After
Spinal cord P4
C
DIC
VR retrograde filling
Texas Red
Fluorescein Dextran
D
DIC
Texas Red
Fluorescein Dextr.
After VR retrograde fill
E
Fluorescein Dextran
ChAT
Merge
L2
F
ChAT+/ Dextran+ MNs (%)
G
Dextran+/ChAT+ MNs (%)
H
P4
P14
P42
L1
ChAT
I
### MNs / section
0.0399
J
### sections used
K
### MNs
Approach 2
Age (d)

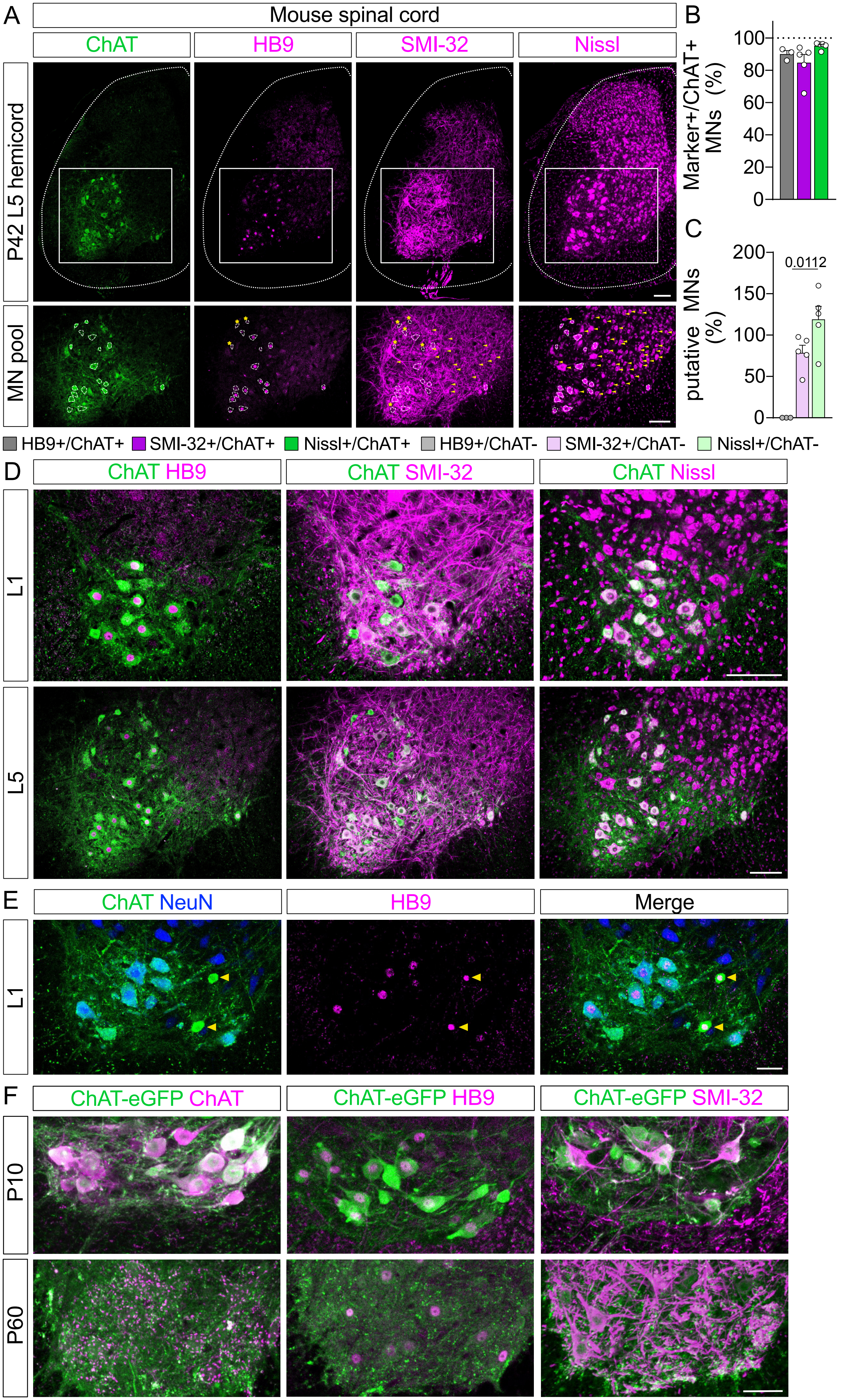


Suppl. Figure 3

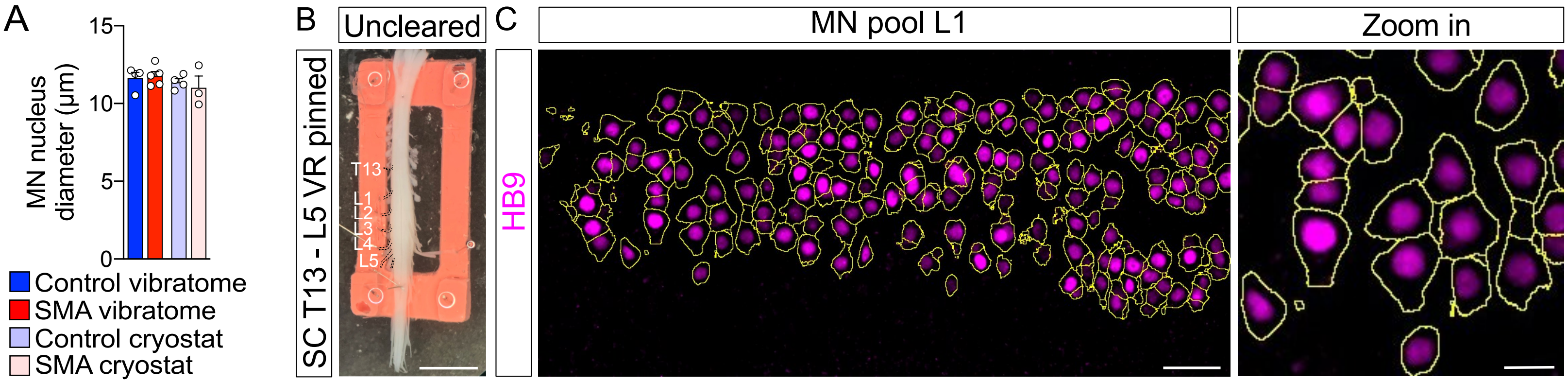
A
15
10
5
0
MN nucleus diameter (µm)
Control vibratome
SMA vibratome
Control cryostat
SMA cryostat
B
Uncleared
SC T13 - L5 VR pinned
T13
L1
L2
L3
L4
L5
C
HB9
MN pool L1
Zoom in

A

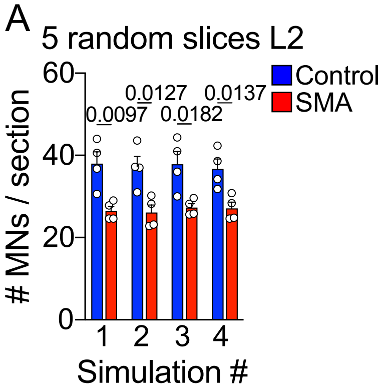
5 random slices L2
Control
SMA
0.0097
0.0127
0.0182
0.0137
### MNs / section
60
40
20
0
1
2
3
4
Simulation #

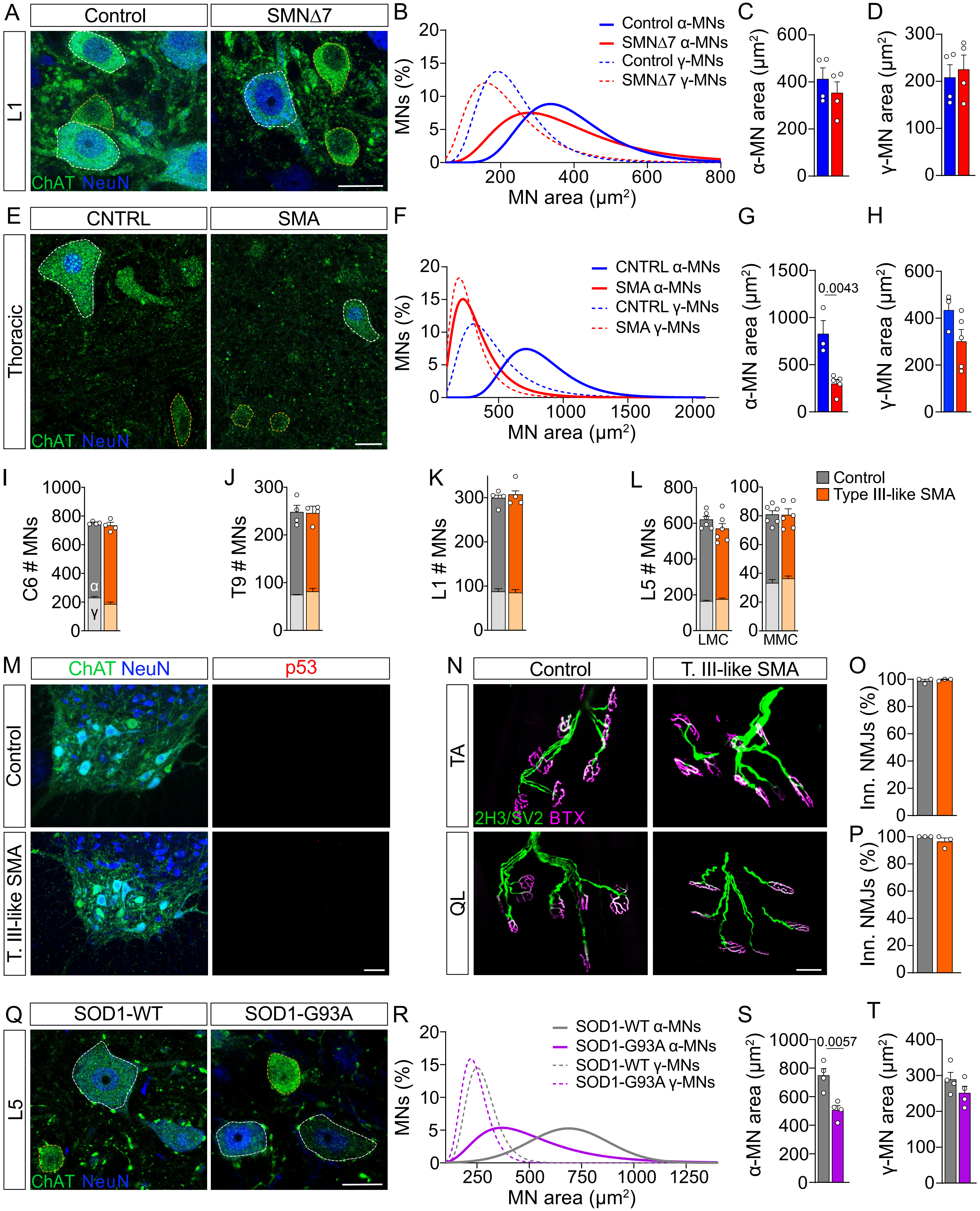
A
Control
SMNΔ7
L1
ChAT NeuN
B
MNs (%)
MN area (µm²)
Control α-MNs
SMNΔ7 α-MNs
Control γ-MNs
SMNΔ7 γ-MNs
C
α-MN area (µm²)
D
γ-MN area (µm²)
E
CNTRL
SMA
Thoracic
F
CNTRL α-MNs
SMA α-MNs
CNTRL γ-MNs
SMA γ-MNs
G
0.0043
H
I
C6 # MNs
α
γ
J
T9 # MNs
K
L1 # MNs
L
L5 # MNs
LMC
MMC
Control
Type III-like SMA
M
p53
T. III-like SMA
N
T. III-like SMA
TA
QL
2H3/SV2 BTX
O
Inn. NMJs (%)
P
Q
SOD1-WT
SOD1-G93A
L5
R
SOD1-WT α-MNs
SOD1-G93A α-MNs
SOD1-WT γ-MNs
SOD1-G93A γ-MNs
S
0.0057
T

**Table S1:** PubMed Search Strategy

| Step | Terms |
|---|---|
| 1 | (atrophy, spinal muscular[MeSH Terms]) OR<br>(spinal muscular atrophy[Title/Abstract]) AND<br>((2005/1/1:2024/2/20[pdat]) AND (english[Filter])) |
| 2 | (murine[Title/Abstract] OR mouse[Title/Abstract] OR<br>mice[Title/Abstract] OR rodent*[Title/Abstract]) AND<br>((2005/1/1: 2024/2/20 [pdat]) AND (english[Filter])) |
| 3 | 1+2 |
| 4 | (((((((SMNΔ7) OR (Δ7)) OR (delta-7)) OR (SMND7)) OR<br>(Smn2B/-)) OR (Taiwanese)) OR<br>“SMA mouse model”[Title/Abstract]) OR<br>“SMA mice”[Title/Abstract] AND (2005/1/1: 2024/2/20 [pdat]) AND<br>(english[Filter]) |
| 5 | 3+4 |

**Table S2**: PICOS Criteria

| Criteria | Inclusion | Exclusion |
| --- | --- | --- |
| **Population** | SMNΔ7: Background must be either Jackson 5025 (FVB.Cg-*Grm7*$^{Tg(SMN2)89Ahmb}$ *Smn1*$^{tm1Msd}$ Tg (SMN2*Δ7) 4299Ahmb/J or state a genetic background of Smn/; SMN2/SMN2; SMNΔ7/ SMNΔ7. The age is between or including P7 and P14.<br><br>Smn$^{2B/-}$: Background must show either a *Smn*$^{2B/2B}$ × *Smn*$^{+/-}$ or state reception from Kothary Lab. The age is between P10 and P28.<br><br>Taiwanese: Background contains 2 copies of SMN gene with a genetic background of Jackson 5058 (FVB.Cg-*Smn1*$^{tm1Hung}$ Tg(SMN2)2Hung/J) or *Smn* $^{-/-}$ *; SMN2* $^{tg/0}$. The age of the two copy model is between P7 and P14. | All mouse models not matching this criteria or altered genetically/cellularly in any further manner. |
| **Intervention** | Studies provide data for a non treated experimental mouse model. | Studies provide no data for a non treated experimental mouse model. |
| **Comparison** | Studies provide data for a non treated control mouse model. | Studies provide no data for a non treated control mouse model. |
| **Outcomes** | MN counts from the anterior spinal cord are reported or a percentage of loss is described. | MN counts in cell culture or not within the spinal cord anterior horn. |
| **Study Design and Publication Type** | Four-arm parallel-group randomized control studies and comparative studies were evaluated. | All other study types. |

**Table S3**: PRISMA Diagram

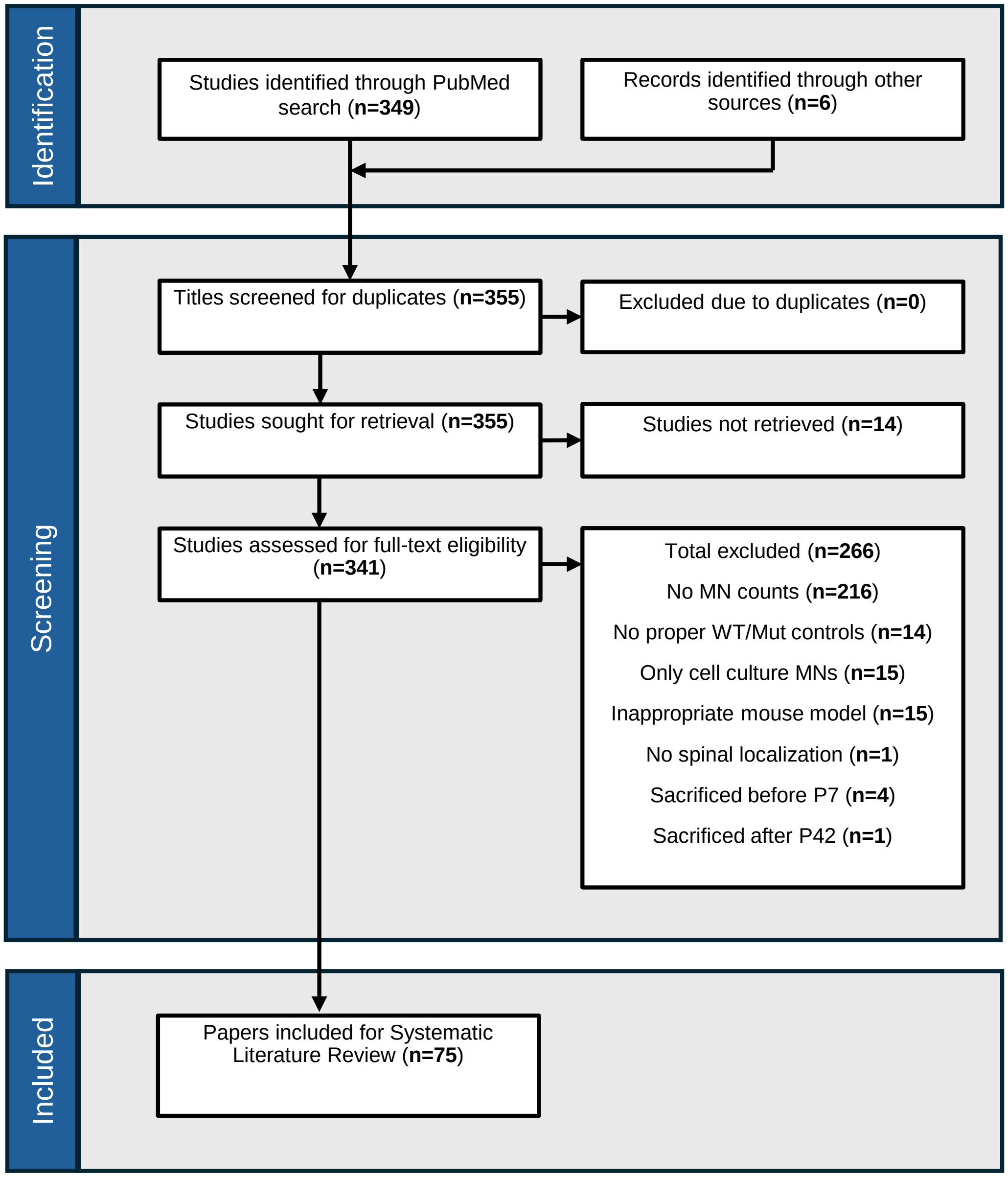

**Table S4**: List of patient tissue used for immunohistochemical analysis

| Control or SMA Type | Age | PMI | Analyzed properties |
|---|---|---|---|
| Control #1 | 5 years | 21 h | MN marker |
| Control #2 | 3 years | ~2 h | MN size |
| Control #3 | 14 years | 2 h | MN size,<br>MN marker |
| Control #4 | 9 months | 14-18 h | MN size,<br>MN marker |
| SMA Type 1 #1 | 14.8 months | < 2 h | MN size |
| SMA Type 1 #2 | 6 years | 2 h | MN size |
| SMA Type 1 #3 | 8 months | 3-9 h | MN size |
| SMA Type 1 #4 | 24 months | 2 h | MN size |
| SMA Type 1 #5 | 11 months | 19 h | MN size |

**Table S5**: List of primary antibodies

| Name | Company | Cat # | Host | Dilution | Labeled Structure |
|---|---|---|---|---|---|
| Bungarotoxin | Invitrogen | B35451 | N/A | 1:500 | NMJ (post-synaptic) |
| Calbindin | Synaptic Systems | 214006 | Chicken | 1:500 | Renshaw cells |
| ChAT | Millipore | AB144P | Goat | 1:500 | MNs |
| FoxP2 | Abcam | ab16046 | Rabbit | 1:5000 | V1 interneurons (nucleus) |
| GFP | Abcam | ab13970 | Chicken | 1:2000 | GFP+ MNs |
| HB9 | Custom made (Wichterle Lab) | | Guinea pig | 1:5000 | MNs (nucleus) |
| Nissl NeuroTrace 500/525 | Invitrogen | N21480 | N/A | 1:500 | MNs |
| NeuN | Millipore | MAB377 | Mouse | 1:2000 | $\alpha$-MNs |
| Neurofilament | DSHB | 2H3, concentrate | Mouse | 1:500 | NMJ (pre-synaptic) |
| Neurofilament-A647 (SMI-32) | Biolegend | 801712 | Mouse | 1:500 | MNs |
| p53 | Leica | NCL-Lp53-CM5p | Rabbit | 1:1000 | Nucleus |
| SV2 | DSHB | SV2, concentrate | Mouse | 1:500 | NMJ (pre-synaptic) |
